# ABLE: Automated Brain Lines Extraction Based on Laplacian Surface Collapse

**DOI:** 10.1101/2022.01.18.476370

**Authors:** Alberto Fernández-Pena, Daniel Martín de Blas, Francisco J. Navas-Sánchez, Luis Marcos-Vidal, Pedro M. Gordaliza, Javier Santonja, Joost Janssen, Susanna Carmona, Manuel Desco, Yasser Alemán-Gómez

## Abstract

The archetypical folded shape of the human cortex has been a long-standing topic for neuroscientific research. Nevertheless, the accurate neuroanatomical segmentation of sulci remains a challenge. Part of the problem is the uncertainty of where a sulcus transitions into a gyrus and vice versa. This problem can be avoided by focusing on sulcal fundi and gyral crowns which represent the topological opposites of cortical folding. We present Automated Brain Lines Extraction (ABLE), a method based on Laplacian surface collapse to segment sulcal fundi and gyral crown lines reliably. ABLE is built to work on standard FreeSurfer outputs, and eludes the delineation of anastomotic sulci while maintaining sulcal fundi lines that traverse the regions with the highest depth and curvature. First, it segments the cortex into gyral and sulcal surfaces; then, each surface is spatially filtered. A Laplacian-collapse-based algorithm is then applied to obtain a thinned representation of the surfaces. This surface is then used for careful detection of the endpoints of the lines. Finally, sulcal fundi and gyral crown lines are obtained by eroding the surfaces while preserving the connectivity between the endpoints. The method is validated by comparing ABLE with three other sulcal extraction methods using the Human Connectome Project (HCP) test-retest database to assess the reproducibility of the different tools. The results confirm ABLE as a reliable method to obtain sulcal lines with an accurate representation of the sulcal topology while ignoring anastomotic branches and the overestimation of the sulcal fundi lines. ABLE is publicly available via https://github.com/HGGM-LIM/ABLE.

## Introduction

The human brain cortex presents a characteristic morphology of folds and fissures. During prenatal and early postnatal neurodevelopment, the process of folding causes a series of sulci and gyri in the cortex. Sulci are defined as the regions of the cortex folded inwards, and gyri as the regions folded outwards^1^. The sulcal regions with high curvature and maximum depth are called sulcal fundi, while the gyral regions with minimum depth values are called gyral crowns.

Accurate in vivo labeling of sulcal fundi and gyral crowns is of interest as accurate fundi/crown labels can be used as landmarks for the deformation fields in brain-surface warping algorithms^2^. Sulcal fundi and gyral crowns also tap topological differences in cytoarchitecture^1^. As such, precise segmentation of fundi/crowns can have important implications for gyral- and sulcal-specific research of brain morphogenesis^3,4^, morphometry^5,6^, the genetics of cortical gyrification^7–9^, cognition^10^, mental illnesses^11^ or neurodegenerative disorders^12,13^.

There is no consensus about the exact location of sulcal fundi and gyral crowns, making the evaluation and assessment of these extraction methods more complex. In addition, the large individual anatomical variability of the human cortex further provides a further challenge for a line extraction process. Nevertheless, automatic line extraction is preferable over manual labeling of sulcal fundi, and gyral crown lines as manual labeling is a very time-consuming task with a poor interrater reliability^14,15^. Initially, sulcal fundi and gyral crowns delineation methods were based on volume-based extraction procedures^16-18^; nowadays, most methods use a surface mesh extracted from MRI volumes, and sulcal fundi/gyral crowns are defined as lines consisting of connected vertices over the mesh. Accurate extraction of the cortical surface mesh is critical for the extraction of sulcal fundi/gyral crowns. Mesh inaccuracies translate into errors in the derived vertex-wise curvature and depth maps, which are essential for line extraction methods. Therefore, the line extraction process relies on the correct outcome of tools such as FreeSurfer^19^, BrainSuite^20^, and BrainVISA’s Morphologist^21^ which allow for the generation of a cortical surface mesh.

There are different approaches for the delineation of sulcal fundi lines from cortical surfaces. Sulcal lines have been defined as the shortest path between two points of the same sulcal basin. This shortest path can be defined in terms of the local cortical surface curvature^22^, or the surface convexity^23^. The shortest-path approach forces sulcal fundi lines to traverse sulcal regions of high curvature or convexity. Other approaches, such as skeleton-based methods, provide skeletonized sulcal lines by pruning sulcal or fold segmentations through the removal of vertices with the lowest curvature^24^, depth^25^, or distance to the fold boundary^26^. The pruning is performed while maintaining the sulcal topology and shape. Both approaches present limitations. For instance, in the shortest-path delineation method, sulcal regions are typically not represented when the sulcus encircles a gyral wall. In addition, the shortest-path and skeleton-based methods rely on the selection of sulci endpoints, i.e., the vertices where a line should start or end, to delineate sulcal fundi lines. The definition of sulcal endpoints is challenging due to the highly variable geometric patterns on the cortical surfaces leading to widely branched sulci, thus making it hard to decide whether the branch of a sulcal fundi/gyral crown line is principal or anastomotic^27^.

To solve this problem, some methods extract the endpoints directly during the sulcal eroding step^26^ or using adaptive thresholding of the Geodesic Path Density Maps (GPDM)^22^. In GPDM, the probability of belonging to a sulcal fundi line is assigned to each vertex on the cortical surface by estimating the “shortest” paths between pairs of vertices in the sulcal basin boundary. This probability is then thresholded to obtain the sulcal fundi lines. This approach leads to sulcal fundi lines traversing the zones with high curvature and depth, but inaccurate in terms of sulcal length^22^. Furthermore, the obtained lines do not represent the entire sulcal topology, especially in bifurcations. The majority of currently available methods contain an explicit endpoint detection step. For instance, in Mindboggle^25^ the endpoints are detected by propagating maximal depth tracks from vertices with median depth towards the sulcal boundary. In Kao et al. (2007)^24^, endpoints are selected as the points of the sulcal boundary that are in the extremes when doing a principal component analysis on a neighborhood of the point. In Topological Graph Representation for Automatic Sulcal Curve Extraction (TRACE)^28^, high curvature sulcal fundi vertices are selected, and the endpoints are defined as the points in the extremes of that set of vertices. This definition leads to the delineation of anastomotic sulci that increase the morphological variability and complicate phenotypic analyses^29^.

In general, progress in the field of line extraction may have also been hampered by the limited availability of source code and a sometimes challenging implementation, i.e., difficult to use. This has resulted in few applications of line extraction methods as well as complicating evaluations of reproducibility. To the best of our knowledge, only Mindboggle (https://mindboggle.info/), TRACE (https://github.com/ilwoolyu/CurveExtraction) and GPDM (https://brainvisa.info/web/) have made their code public and easily are available.

This work presents a new method named **A**utomated **B**rain **L**ines **E**xtraction Based on Laplacian Surface Collapse (**ABLE**) to reliably and automatically segment sulcal fundi and gyral crown lines. This algorithm is implemented in a publicly available MATLAB toolbox (https://github.com/HGGM-LIM/ABLE), and it was developed to work over the standard FreeSurfer outputs (cortical surfaces and curvature maps). It is also compatible with other cortical surface extraction software suites as long as these provide a surface mesh and a vertex-wise curvature map.

Our proposed methodology overcomes limitations such as the under- or overestimation of the sulcal length and the spilling of sulcal lines into gyral regions. ABLE also eludes the tracing of anastomotic sulci^27^ while maintaining the lines traversing through regions with high curvature values inside the same sulcal basin. Besides, it extends the existing skeleton-based methods by filtering the sulcal basins, applying Laplacian collapse, and detecting the endpoints of the resulting skeletonized mesh. A test-retest dataset is used for assessing and validating the results by comparing ABLE with the three publicly available methodologies: BrainVISA’s GPDM^22^, Mindboggle^25^, and TRACE^28^.

## Results

### Brain lines: gyral crowns and sulcal fundi

Figure 1 displays the brain lines obtained by using the developed method, ABLE. The images in Figure 1A show how the gyral lines traverse the top of the gyral regions. Figure 1B depicts the sulcal lines over the inflated surface along with the gyral crown; the different sulcal basins are represented with different colors. The zoom-in of these images compares the lines obtained by ABLE (red) to the ones extracted by using GPDM (blue), Mindboggle (orange) and TRACE (green).

**Figure 1.**
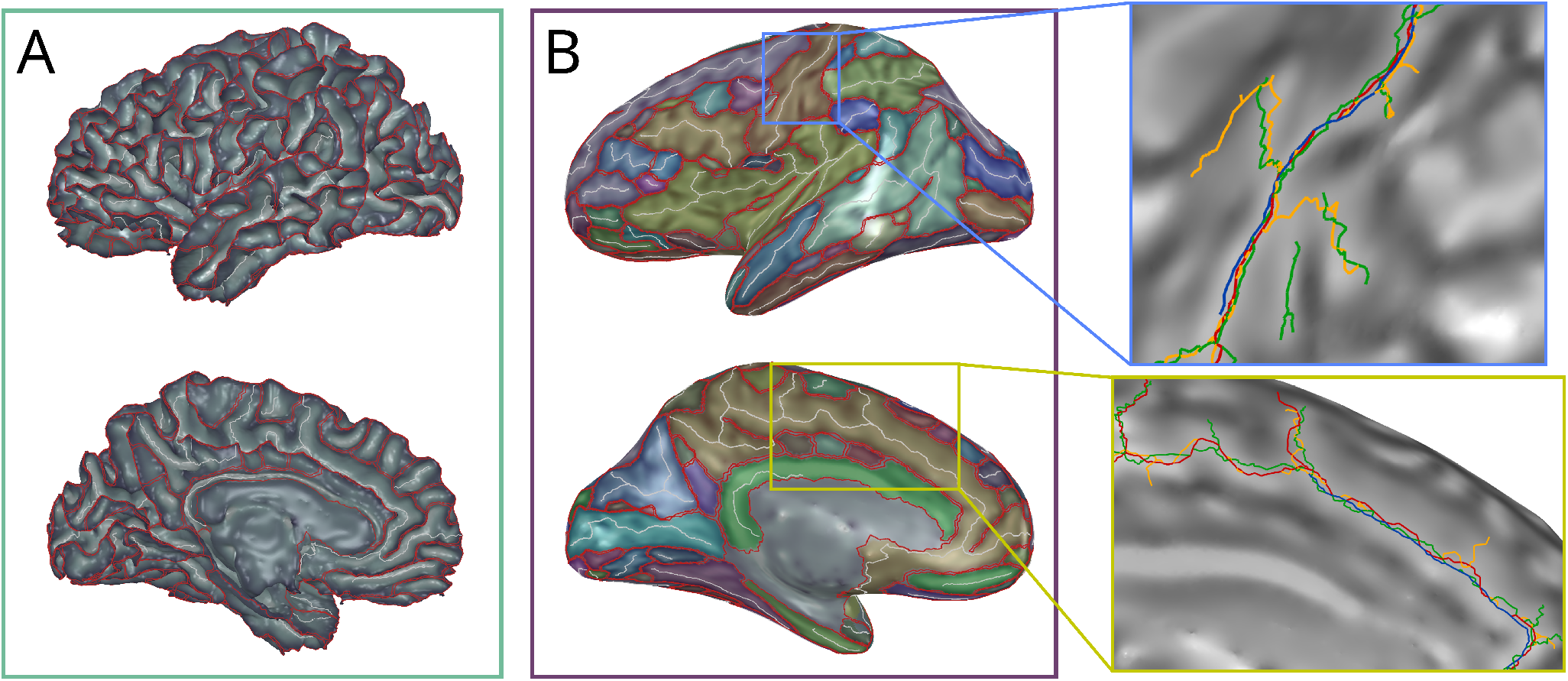
Individual brain lines computed using ABLE. (A) Gyral crowns (red) and sulcal fundi (white) lines depicted on the white matter surface. The greyscale map represents the curvature values, where higher curvature values (fundi) are lighter and lower curvature values (gyri) are represented in dark. (B) shows the gyral crowns (red) and sulcal fundi (white) depicted on the inflated white surface. Superimposed to the curvature map is the subdivision of the different sulcal basins obtained by our method. Two zoom-in images show the lines for central and cingulate sulci obtained by the different sulcal lines extraction methods. **Note:** Lines in blue correspond to GPDM, red to ABLE, green to TRACE, and orange to Mindboggle.

### Sulcal fundi curvature and depth

The analysis of the differences in sulcal fundi curvature (Figure 2) shows that ABLE presents significantly higher curvature values than the other methods except for GPDM (*q* < 0.05, see Supplementary Table S1). ABLE and GPDM are very similar in terms of mean curvature values in all the sulci, except for the right calcarine sulcus (*t* = −4.06, *q* = 0.002) and parieto-occipital fissure (*t* = −3.12, *q* = 0.031), where GPDM scores higher curvatures; and the right cingulate sulcus (*t* = 6.57, *q* < 0.01), with ABLE producing higher curvature than GPDM.

**Figure 2.**
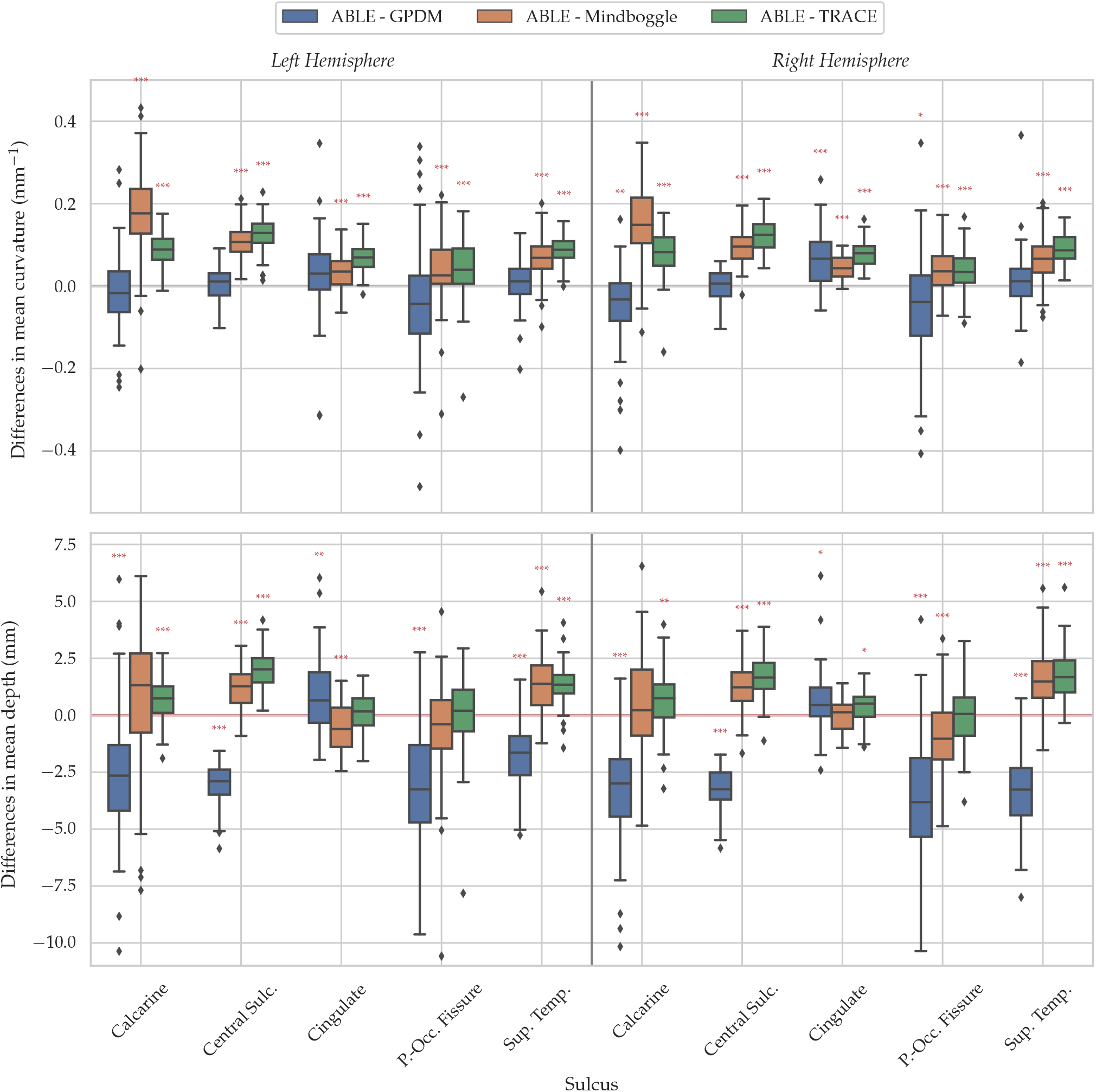
Differences in curvature (top) and depth (bottom) between ABLE and the other reference methods. Symbols “*”, and “***”, stand for significant differences between methods with FDR-corrected p-values lower than 0.05, 0.01, and 0.001 respectively.

In case of sulcal fundi depth differences (Figure 2 and Supplementary Table S1), ABLE shows lower depth values than GPDM in every sulcus (with t-values ranging from −27.55 to −7.77, and *q* < 0.001) except for the left and right cingulate sulcus (with *t* = 4.25, *q* = 0.001; and *t* = 3.24, *q* < 0.001 respectively). ABLE shows larger depth values compared to Mindboggle for the central sulcus (left: *t* = 12.34, *q* < 0.001 and right: *t* = 9.46, *q* < 0.001) and superior temporal sulcus (left: *t* = 8.54, *q* < 0.001; and right:*t* = 9.58, *q* < 0.001) in both hemispheres. It also has lower values in the left cingulate sulcus (*t* = −4.70, *q* < 0.001) and right parieto-occipital fissure (*t* = −4.42, *q* < 0.001). No other significant differences are found (*q* < 0.05). Finally, ABLE has higher depth values than TRACE in all the sulci except for the left cingulate sulcus and bilateral parieto-occipital fissures where no significant differences are found.

### Sulcal fundi longitudes

Figure 3 shows, for the selected sulci, the regression results between the sulcal fundi longitudes computed for each method (y-axes) versus the reference values, BrainVISA’s Morphologist, (x-axes).

**Figure 3.**
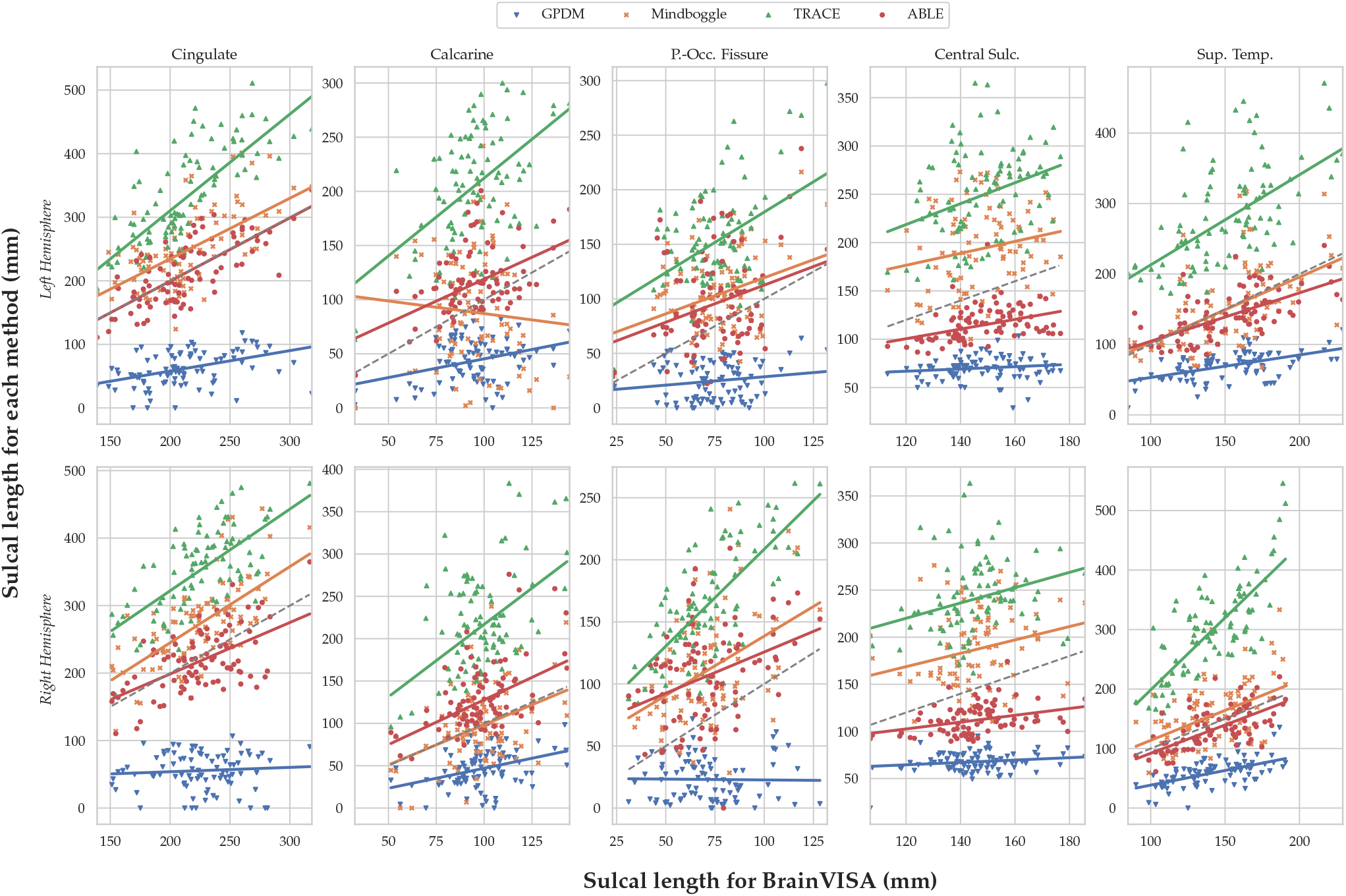
Scatter plots and regression of the longitudes obtained using the different sulcal lines extraction methods and the BrainVISA longitudes for the studied sulci. The dashed line the regression coefficient = 1 and intercept = 0.

In Table 1 we provide the coefficient of determination (*R*^2^), the slope and the associated T-value for each linear regressions analysis. In Table S4 of the Supplementary material we provide also the results for the regressions intercepts. We can observe from these values that regressions between ABLE and BrainVISA longitudes present slopes, in average, closer to one (ranging between 0.65 and 1.20) except for the central sulcus, where values are lower (0.49 for left and 0.38 for right). Every coefficient value for ABLE is significant (*q* < 0.05). For TRACE longitudes, the obtained slopes (range: 1.05 – 2.39) and intercepts (range: 12.27 – 154.23) are consistently over one and zero respectively, except for the right central sulcus (slope = 0.61, *q* = 0.190) and right superior temporal (intercept = −36.53). Mindboggle values also presents slopes near one for all the sulci (range: 0.68 – 1.05), except for the left calcarine (slope = −0.01, *q* = 1.000). GPDM presents flatter slopes with values lower than one (range: 0.56 – 0.28 where *q* < 0.05) for all the sulci. Regarding the obtained *R*^2^ values, these are quite similar for ABLE and TRACE (ABLE is between 0.08 and 0.52, while TRACE shows values between 0.10 and 0.58).

**Table 1.** Coefficient of determination, slope coefficients and associated T-values for the regressions of the longitudes obtained by the different methods and the reference values (i.e., BrainVISA).

| Sulcus Name | Longitude regression |  |  |  |  |  |  |  |  |  |  |  |
| --- | --- | --- | --- | --- | --- | --- | --- | --- | --- | --- | --- | --- |
| | $R^2$ <sup>a</sup>($p$-FDR<sup>b</sup>) | ABLE<br>Coefficient<sup>c</sup> | $T^*$ | $R^2$ ($p$-FDR) | GPDM<br>Coefficient | $T$ | $R^2$ ($p$-FDR) | TRACE<br>Coefficient | $T$ | $R^2$ ($p$-FDR) | Mindboggle<br>Coefficient | $T$ |
| **Left Hemisphere** |  |  |  |  |  |  |  |  |  |  |  |  |
| Calcarine | 0.24 (**<0.001**) | 0.82 (0.52, 1.12) | 5.41 | 0.10 (**0.010**) | 0.33 (0.13, 0.53) | 3.22 | 0.25 (**<0.001**) | 1.40 (0.89, 1.90) | 5.51 | -0.01 (1.000) | -0.10 (-0.68, 0.48) | -0.35 |
| Central Sulcus | 0.15 (**<0.001**) | 0.49 (0.25, 0.73) | 4.07 | 0.00 (1.000) | 0.10 (-0.08, 0.27) | 1.09 | 0.10 (**0.009**) | 1.05 (0.41, 1.70) | 3.26 | 0.03 (0.311) | 0.60 (-0.03, 1.23) | 1.88 |
| Cingulate | 0.52 (**<0.001**) | 0.97 (0.78, 1.17) | 9.78 | 0.14 (**0.001**) | 0.28 (0.14, 0.42) | 3.97 | 0.56 (**<0.001**) | 1.51 (1.23, 1.79) | 10.65 | 0.39 (**<0.001**) | 1.01 (0.74, 1.27) | 7.56 |
| Parieto-Occipital Fissure | 0.08 (**0.02**) | 0.71 (0.24, 1.19) | 2.98 | 0.02 (0.558) | 0.15 (-0.04, 0.35) | 1.58 | 0.23 (**<0.001**) | 1.22 (0.76, 1.68) | 5.29 | 0.12 (**0.004**) | 0.68 (0.30, 1.06) | 3.57 |
| Superior Temporal | 0.41 (**<0.001**) | 0.68 (0.51, 0.85) | 7.95 | 0.28 (**<0.001**) | 0.33 (0.22, 0.44) | 6.01 | 0.33 (**<0.001**) | 1.27 (0.89, 1.65) | 6.65 | 0.26 (**<0.001**) | 0.92 (0.60, 1.24) | 5.66 |
| **Right Hemisphere** |  |  |  |  |  |  |  |  |  |  |  |  |
| Calcarine | 0.26 (**<0.001**) | 1.20 (0.78, 1.61) | 5.74 | 0.13 (**0.002**) | 0.46 (0.22, 0.70) | 3.82 | 0.24 (**<0.001**) | 1.71 (1.08, 2.34) | 5.43 | 0.16 (**<0.001**) | 1.01 (0.54, 1.48) | 4.28 |
| Central Sulcus | 0.11 (**0.01**) | 0.38 (0.16, 0.59) | 3.48 | 0.05 (0.111) | 0.18 (0.03, 0.33) | 2.36 | 0.04 (0.190) | 0.61 (0.04, 1.18) | 2.12 | 0.03 (0.253) | 0.58 (-0.00, 1.17) | 1.99 |
| Cingulate | 0.32 (**<0.001**) | 0.78 (0.54, 1.01) | 6.60 | -0.01 (1.000) | 0.04 (-0.13, 0.21) | 0.47 | 0.44 (**<0.001**) | 1.23 (0.94, 1.52) | 8.41 | 0.45 (**<0.001**) | 1.22 (0.94, 1.51) | 8.60 |
| Parieto-Occipital Fissure | 0.11 (**0.01**) | 0.65 (0.27, 1.02) | 3.46 | -0.01 (1.000) | -0.00 (-0.18, 0.17) | -0.05 | 0.50 (**<0.001**) | 1.55 (1.22, 1.87) | 9.44 | 0.21 (**<0.001**) | 0.99 (0.59, 1.39) | 4.91 |
| Superior Temporal | 0.42 (**<0.001**) | 0.91 (0.69, 1.14) | 8.14 | 0.39 (**<0.001**) | 0.56 (0.41, 0.71) | 7.59 | 0.58 (**<0.001**) | 2.39 (1.96, 2.82) | 11.17 | 0.24 (**<0.001**) | 1.05 (0.67, 1.44) | 5.44 |

<sup>a</sup>The comparisons between the groups were conducted using pairwise $t$ -tests.
<sup>b</sup> $p$ -values in bold remain significant after entering all uncorrected $p$ -values ( $N = 120$ ) into a False Discovery Rate (FDR) correction for multiple comparisons with $q = 0.05$ .
<sup>c</sup>CI = 95% confidence interval.

Figure 4 shows Bland-Altman plots comparing the length obtained using the line extraction methods against the BrainVISA length for all the sulci. The plot shows that the mean length difference between ABLE and BrainVISA is the lowest among all the sulcal lines extraction methods (2.24 mm). In the case of GPDM, there is an underestimation of the mean sulcal length (−84.66 mm) compared to the mean length reported by BrainVISA. Finally, Mindboggle and TRACE depict mean length differences of 25.89 mm and 113.8 mm respectively, compared to the reference values (i.e., BrainVISA).

**Figure 4.**
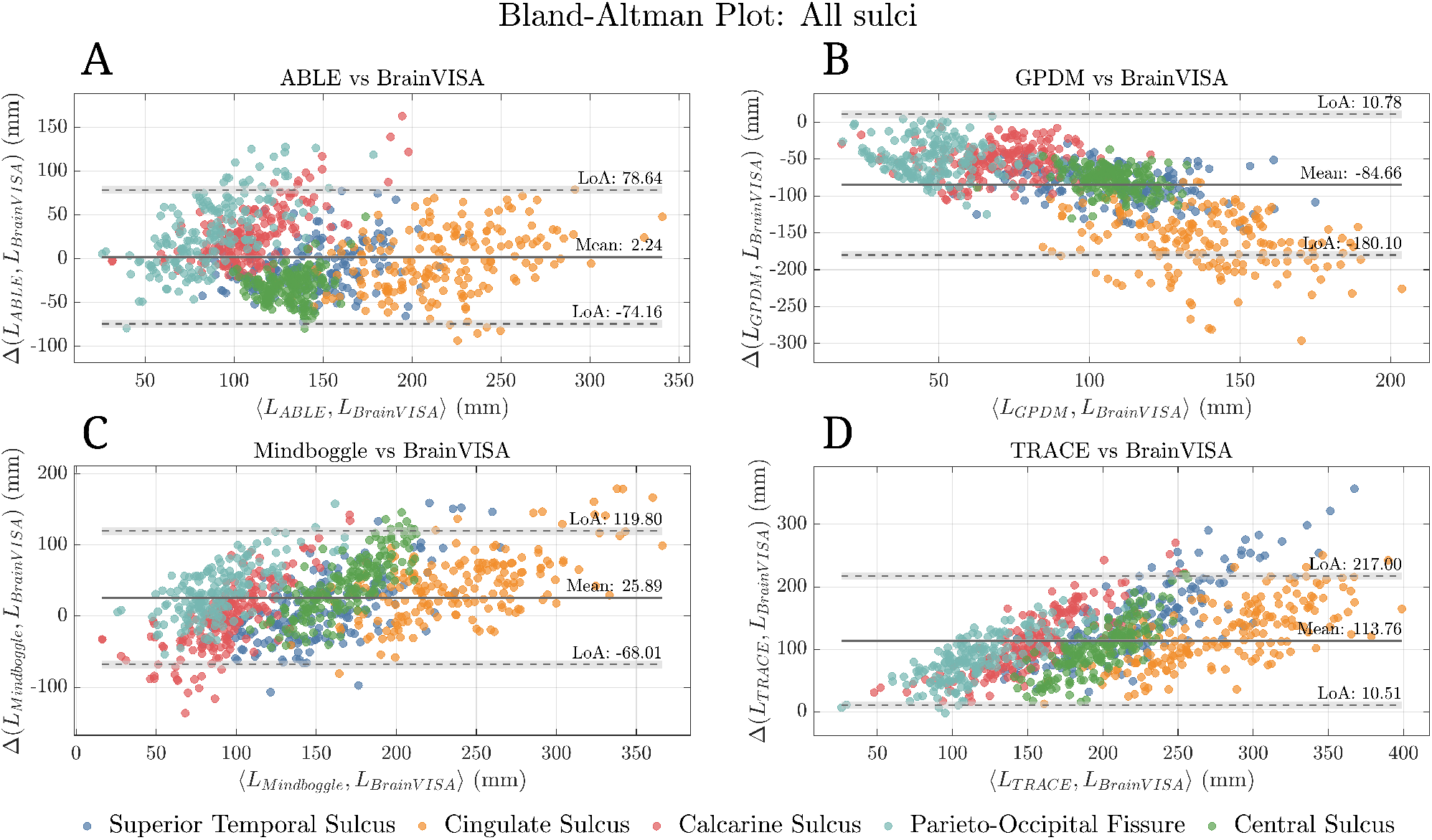
Bland-Altman plots comparing the four methods (ABLE, GPDM, Mindboggle, and TRACE) against BrainVISA sulcal lengths. The plot includes every sulcus of interest (each one in a different color) in both left and right hemispheres. Limits of Agreement (LoA) are estimated as the mean difference ±1.96 standard deviation.

More information about the mean length differences between each method and BrainVISA for each individual sulcus can be found in Table S3 and in the Bland-Altman plots shown in the Figures S1-S10 of the Supplementary material.

### Reproducibility: Hausdorff distances between test and retest

The logarithms of the mean and maximum Hausdorff distances are used to assess the reproducibility of each method. These logarithms for every sulcus are depicted in Figure 5 and summarized in Supplementary Table S2 along with the statistical results. TRACE presents the lowest averages (−0.16 – −0.04) and standard deviations of the mean Hausdorff distances for most of the sulci, followed by ABLE (mean values between −0.18 and 0.10). ABLE performs better than Mindboggle for the bilateral calcarine (left: *t* = −4.81, *q* < 0.001, right: *t* = −4.63, *q* < 0.001), bilateral central (left: *t* = −7.40, *q* < 0.001, right: *t* = −6.37, *q* < 0.001), and left superior temporal sulcus (*t* = −5.66, *q* < 0.001). The significant differences with GPDM are limited to the left parieto-occipital fissure (*t* = −4.04, *q* = 0.003). For the maximum Hausdorff distances, significant differences between ABLE and TRACE are limited to the left calcarine sulcus (*t* = 3.42, *q* = 0.014) and right central sulcus (*t* = −4.33, *q* = 0.001).

**Figure 5.**
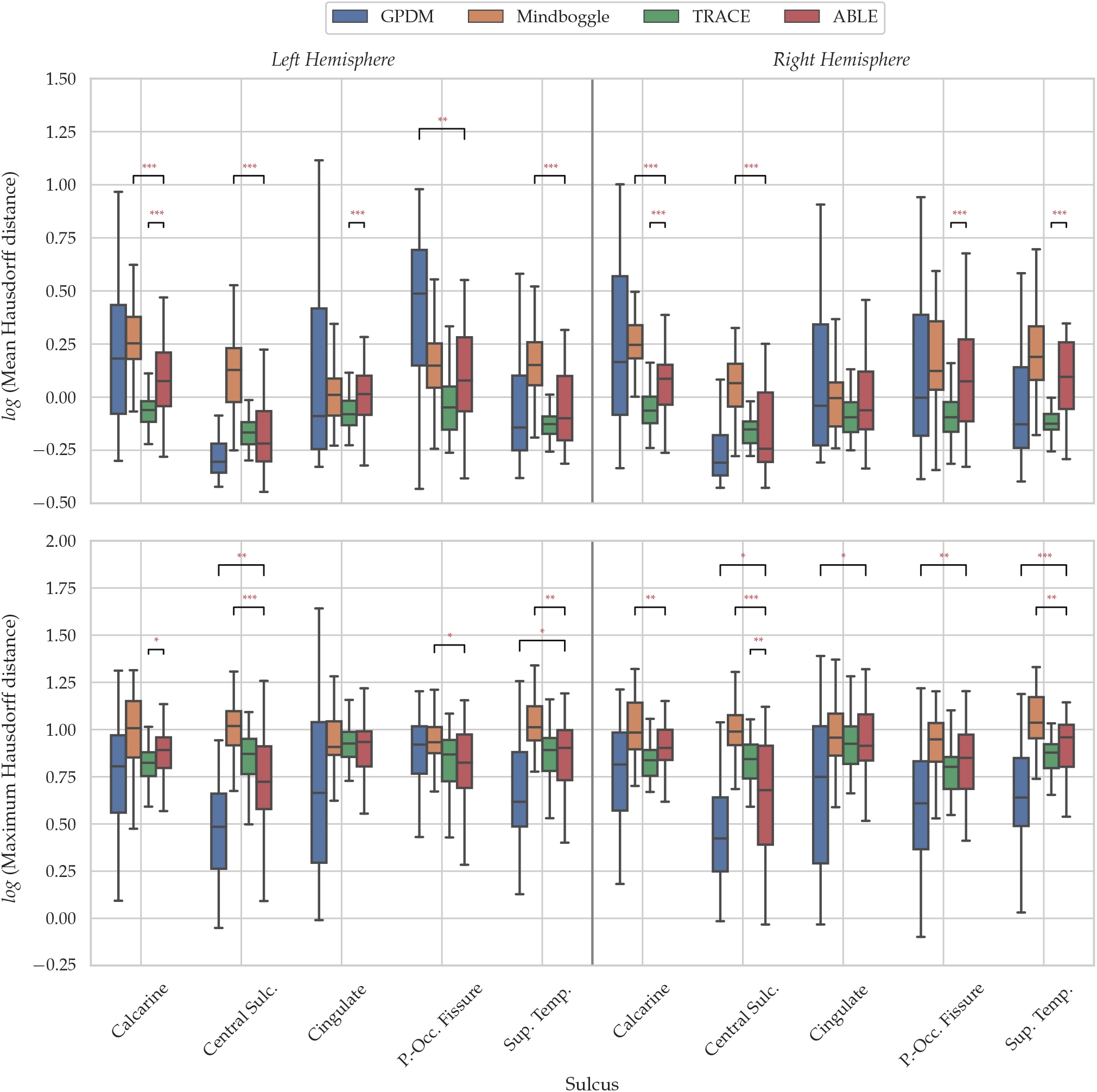
Comparisons of logarithms of mean (top) and maximum (bottom) Hausdorff distances between ABLE and the reference methods. These distances are computed between sucal lines coordinates extracted for test and retest acquisitions. Symbols “*”, “**”, and “***”, stand for significant differences from a t-test with FDR-corrected p-values lower than 0.05, 0.01, and 0.001, respectively.

## Discussion

We present a new method for the automated delineation of sulcal fundi and gyral crown lines and validate our approach against three publicly available methods. Globally, our approach performs similarly or better for the tested metrics, providing a reliable and improved method for extracting brain sulcal fundi lines.

The lack of a unique criterion for the definition of sulcal fundi lines makes it difficult to judge which method is more accurate. Indeed, the sulcal fundi lines should traverse regions with the highest mean curvature for some methods, while for other methods, the lines should traverse regions with the highest depth. Depth-based methods use either Euclidean or geodesic^25^ definitions of depth or a mixture of them^24^ which further illustrates the lack of a single set of criteria for evaluation. Depending on the objective, one method can be more appropriate than another. For example, GPDM minimizes the labeling of bifurcations at the expense of obtaining shorter sulcal fundi lines with high curvature values while leaving extensive regions of the sulcal basins without lines. On the other side, TRACE and Mindboggle include small sulcal bifurcations in their labeling at the expense of having much lower curvature and depth values along these lines. ABLE is designed to reach an intermediate solution, labeling branches but avoiding anastomotic sulci.

Assuming that sulcal fundi run across paths with the highest depth and curvature, these values are used as quantitative metrics to evaluate the extraction methods. GPDM is the method with the best performance in both metrics, followed by ABLE. These results indicate that these two methods traverse the sulci’s fundus more precisely than the other methods. This result is coherent with what is presented in Figure 1, as GPDM only labels regions with very high curvature located at the center of the sulcal basins. Besides, small bifurcations, perpendicularly to the main sulcal line, tend to rapidly decrease in curvature and depth; both Mindboggle and TRACE tend to include these branches and therefore present lower values.

Similarly, sulcal length comparisons demonstrate that ABLE is the closest to BrainVISA (Figure 4A), except for the Parieto-Occipital Fissure, while GPDM tends to considerably underestimate the length of the sulcal lines (Figure 4B). On the other hand, TRACE’s longitude values are notably overestimated (Figure 4D). The longitudes obtained by Mindboggle are in a range between TRACE and ABLE (Figure 4C). These findings confirm that GPDM produces shorter sulcal lines, while TRACE and Mindboggle label many short branches resulting in greater sulcal length. This effect can be observed in Figure 4, where TRACE and Mindboggle show larger differences for higher mean length values. ABLE establishes a compromise between both alternatives, which is more precise given the results of the comparison against the reference sulcal length values extracted from BrainVISA. However, the sulcal line endpoints could affect the length values of ABLE, TRACE, and Mindboggle. Incorrect detection of endpoints leads to an over- or underestimation of the sulcal length, e.g., due to in- or exclusion of sulcal branches. Endpoint detection relies heavily on the selected parameters (i.e., the filtering iterations and the neighborhood size around the endpoints), and an exhaustive study should be performed to estimate their optimal values. The values employed in this paper were selected after an evaluation using healthy young subjects of different sex and age. These default values can be fine-tuned and adjusted to each particular dataset.

Regarding the robustness and reproducibility, TRACE presents lower values in terms of mean Hausdorff distance, producing in some cases values lower than 1 mm. ABLE performs better than GPDM and Mindboggle for the mean Hausdorff distance, showing mean distance values close to 1 mm. This distance value is equivalent to a sulcal line displacement of one edge over the cortical surface between the lines computed from test and retest datasets. TRACE is the most reliable method in terms of mean Hausdorff distances. However, this result must be interpreted with caution. First, TRACE does not traverse the vertices on the surface; it follows a path through different parts of the edges^28^. This makes the method less dependent on the topology of the mesh in regions where vertices are more scattered. Second, TRACE labels more sulcal line bifurcations compared to the other methods. It is more probable to find a closer distance between sulcal lines from the test and retest when those lines contain more points than lines compounded by fewer points. Maximum Hausdorff distances for the four methods are similar, with few significant differences between ABLE and the other methods. GPDM shows slightly better results than the other methods for this metric, supporting the hypothesis that a higher number of line points improves the performance in mean Hausdorff distance, as substantial errors tend to be masked when more points are counted.

Reliable and reproducible sulcal lines extraction methods open many opportunities for the study, in vivo, of pathologies inducing topological changes in the brain cortex such as Alzheimer’s disease or mental disorders. For example, there is ample post-mortem evidence for supragranular layer damage associated with schizophrenia but in vivo evidence is sparse^30^. Supragranular layers are thicker in sulci, particularly at the fundus, compared to gyri. Therefore, one could hypothesize that disease-associated supragranular layer damage may be more prevalent in sulcal fundi as compared to gyral crowns. By assessing cortical thickness (as a marker of cortical layer damage) at fundi/crowns, we can examine this hypothesis in vivo.

Finally, ABLE relies on gyral crown lines to define the sulcal basins containing the sulcal lines. Due to the lack of available methods for delineating the gyral crown lines, we did not implement a formal evaluation of the extraction of these lines. Future studies should be performed to quantitatively and qualitatively assess the performance of ABLE and other gyral crowns lines extraction.

In summary, it is noteworthy that ABLE is the most balanced method among those compared. For sulcal length, ABLE performs better than GPDM in terms of reproducibility and correlation with BrainVISA sulcal length values, although GPDM provides sulcal lines with a moderately higher curvature or depth. Also, ABLE performs better than TRACE in mean curvature, depth, and correlation with BrainVISA for sulcal length, although TRACE’s mean Hausdorff distance values are slightly better. These findings confirm ABLE as a reliable method to extract sulcal lines with an accurate representation of the sulcal topology while avoiding anastomotic branches and the overestimation of the sulcal fundi lines.

## Conclusion

In this paper, we present a new method for the automatic extraction of sulcal and gyral lines. The method uses smoothed patches of the cortical surfaces to generate collapsed meshes where the endpoints of the sulcal and gyral crown lines are detected. By accurately detecting these endpoints, smaller anastomotic sulcal and gyral crown lines are discarded while preserving the main sulcal and gyral crown line topology. The results demonstrate that lines obtained by ABLE contain high curvature and depth, and their length strongly correlates with the state-of-the-art reference values. This method also delivers correct results in terms of reproducibility (mean and maximum Hausdorff distances). The ABLE algorithm offers a publicly available alternative for the extraction of sulcal and gyral crown lines (https://github.com/HGGM-LIM/ABLE).

## Methods

The proposed method employs the cortical surfaces and their corresponding curvature maps to automatically extract the sulcal fundi and gyral crown lines. These cortical surfaces are triangular meshes defined in the boundaries between grey and white matter (white surface) and grey matter and cerebrospinal fluid (pial surface).

### Geodesic depth map estimation

A geodesic depth map for the pial surface is computed by using the methodology developed in Rettmann et al. (2002)^31^. The depth map computation begins with the estimation of an outer hull surface that allows the identification of the gyral parts of the surface. The outer hull surface is a smooth envelope wrapped around the pial surface that does not encroach into the sulci. The process of generating this outer hull can be seen as the surface equivalent of a topological closing. The points, over the hull surface, intercepting the pial surface are labeled as zero-depth points and taken as gyral regions. Hence, sulcal regions are compound by the remaining pial surface points. Finally, the depth map is calculated as the minimum geodesic distance from the points in the sulcal regions to any of the zero-depth vertices. The geodesic distance was computed using the algorithm proposed in Mitchell et al. (1987)^32^ as implemented in the Geometry Processing Toolbox (gptoolbox)^33^ for MATLAB.

### Gyral crowns lines and sulcal fundi extraction

The computation of sulcal fundi and gyral crown lines consists of several steps (see the complete workflow in Figure 6) that process the white and pial surfaces, along with the curvature and geodesic depth maps, to obtain the lines traversing the gyral crowns and sulcal fundi.

**Figure 6.**
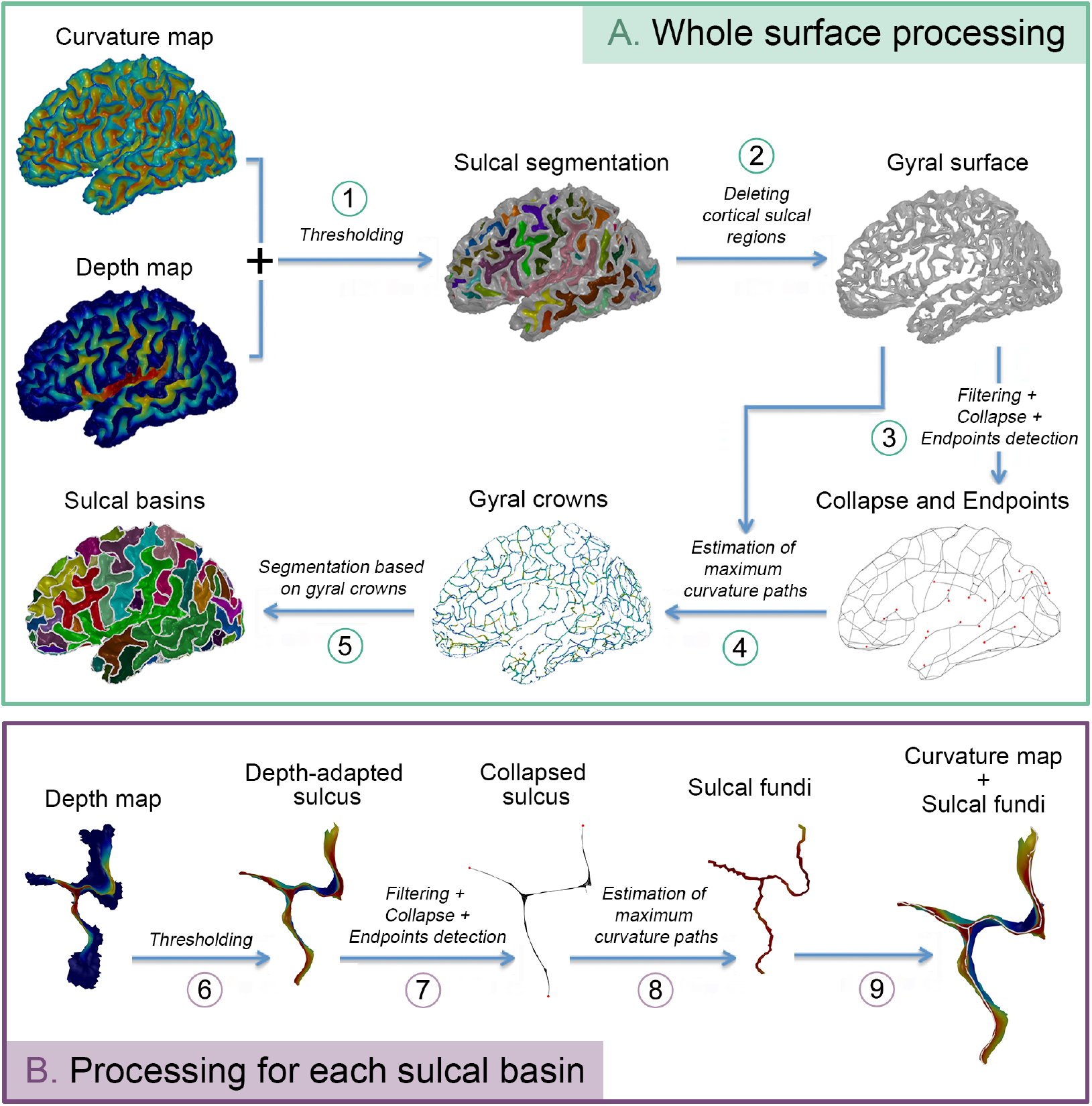
Processing workflow summary. A) Workflow for the extraction of gyral crown lines and the sulcal basins used for the sulcal fundi lines extraction. Curvature and depth maps are thresholded (1) to obtain an initial cortical surface parcellation into sulcal basins and gyral regions (2). These gyral regions are filtered and collapsed; then, an endpoint detection algorithm is applied (3). The gyral crown lines are estimated by decimating the surface while preserving the endpoints and sulcal holes (4). These gyral crown lines are used to delimit the sulcal basins containing the sulcal fundi lines (5). B) Processing steps applied for each sulcal basin. The depth map is thresholded (6) to remove basins sections containing gyral regions, and the resulting surface is filtered and collapsed to detect the sulcal endpoints (7). Finally, as in the gyral crown extraction, the sulcal basin surface is decimated, preserving holes and endpoints (8) to obtain the sulcal fundi lines (9).

#### Sulcal segmentation

The initial step subdivides the white surface into meshes representing the gyral regions and the sulcal basins using the curvature and depth maps. The white surface is segmented, labeling as sulcal regions the vertices with positive curvature and depth over 1mm. Positive curvature is often used as a boundary threshold between sulcal and gyral regions but tends to include more vertices of the sulcal regions than desired due to the positive curvature of the sinuous parts of sulcal banks. These errors are minimized by adding a depth threshold of 1mm. This way, not only the vertices with high curvature are selected, but vertices must also be deep enough to be considered sulcal regions. Thus, deeper regions with curvature values close to zero are included in the sulcal basins. This thresholding splits the white matter mesh into two: sulcal and gyral surfaces (see Figure 6A-1).

#### Gyral crowns extraction

The gyral crown lines are extracted from the gyral surface (Figure 6A-2). White matter surface is used for this step as the gyral regions are more defined, presenting a marked path that is not as noticeable in the pial surface. The generation of these lines needs to fulfill two main conditions: 1) the number and the topology of the holes present in the gyral surface due to the deletion of the sulcal regions must be preserved in the representation of the gyral line, and 2) the gyral morphology should be maintained by preserving the gyral endings. To fulfill the second condition, we must detect and label the gyral endings. The endings are located in regions where the gyral surface loses depth and curvature and converges to a sulcal region. The gyral surface is collapsed to detect the endpoints of the gyral endings (Figure 7A).

**Figure 7.**
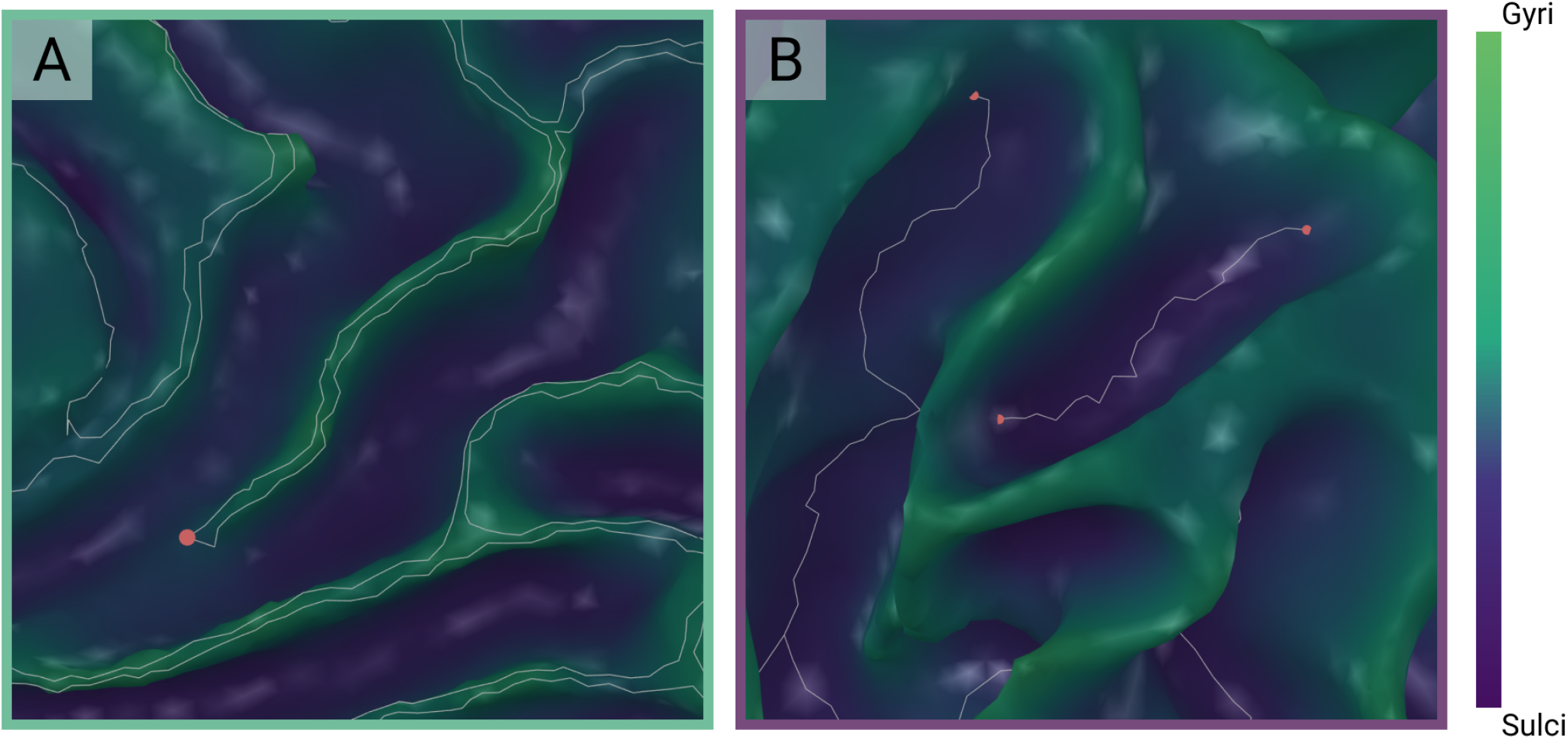
A) Delineation of gyral crown lines (in white) in a gyral branch that fades over the left anterior cingulate sulcus. In red, the estimated endpoint for this protruding gyral region. B) Extracted sulcal lines (white) and endpoints (red) for different sulcal basins located at the left parietal cortex.

Before applying the Laplacian collapse, an iterative mean filtering process is applied to smooth the gyral surface mesh spatially. This way, the appearance of spurious branches in the collapsed surface due to transitional vertices in the sulci-gyri boundary is prevented. In each iteration of the filtering process, the surface vertices’ positions are updated by moving them towards the center of masses of the blocks formed by each vertex and its neighbors. The number of iterations determines the degree of filtering of the surface. For the tested dataset, values between 80 and 120 were, after visual inspection, adequate for obtaining a gyral surface mesh retaining the main shape of the surface and without high-frequency spatial variations. In this work, a value of 100 was selected.

Afterward, Laplacian-collapse algorithm^34^ is applied to the filtered mesh to obtain a thinned representation of the gyral surface. In this step, the thinned surface is obtained by collapsing the vertices of the mesh to a strip by applying an implicit Laplacian smoothing with global positional constraints^34^ while preserving the original connectivity and topology of the mesh. Specifically, an implementation of the algorithm^34^ in gptoolbox^33^ was used for this purpose. Only the geometry contraction step of this implementation is applied, while the connectivity surgery and refinement steps are avoided. The resulting surface is a thinned version of the original gyral surface that maintains the connectivity and topology of the original gyral surface mesh. Finally, the extreme branch points of this surface are detected and labeled as the gyral endpoints (Figure 6A-3).

The detection of the gyral endpoints is based on the method proposed by Kao et al. (2007)^24^. Every vertex in the gyral thinned mesh is evaluated by taking its neighboring vertices and applying a Principal Components Analysis (PCA) to the vertices’ coordinates in order to extract the extreme points along the principal dimension. Gyral endpoints are defined as the points that appear as extreme points in every neighborhood they belong to. For each vertex, the neighboring points are the vertices located at a maximum geodesic distance of 20mm. This threshold value was an appropriate value for the tested dataset to extract the main gyral branches while removing the shortest ones. To accelerate the endpoint detection process, given the vast amount of vertices in the gyral surface, only the vertices forming an angle lower than 30° with their corresponding neighbors in a face are evaluated. This is because only the vertices in the acute “corners” of the surface are suitable to be endpoints, so there is no much interest in evaluating the rest of the points.

The gyral crown lines must connect the endpoints while enclosing the holes in the surface. For this, the gyral surface mesh is eroded during an iterative process in which, in each iteration, the boundary vertex (and its associated faces) are deleted from the mesh. Three main conditions should be fulfilled to eliminate this boundary vertex: 1) the vertex is not an endpoint; 2) its removal does not change the topology of the gyral surface or the endpoints connectivity; 3) it should be the vertex with the minimum curvature value among those satisfying requirements 1 and 2. The conditions are assessed by listing the boundary vertices according to their curvature values and evaluating, starting from the one with the lower curvature, if the previously described conditions are achieved.

The iterations are performed until no other surface vertex is removed (Figure 6A-4). The output of this process is a surface containing the remaining vertices and their shared faces. This surface is simplified to a set of lines by keeping only the external boundary edges. These lines represent the maximum curvature paths connecting the detected gyral surface endpoints and enclosing the surface holes. Finally, the sulcal basins are defined as the individual sulcal surfaces enclosed by these gyral crown lines (Figure 6A-5).

#### Sulcal fundi extraction

The methodology used to obtain the sulcal lines is similar to the one described in the previous section. In this case, the surface used is the pial one, as is the one presenting more defined sulci. The use of one surface or the other is possible due to the fact that FreeSurfer surfaces have a correspondence in the number of vertices and the order of the faces, so different maps and indices on the surface are valid for both of them.

For each sulcal basin extracted from the pial surface, the vertices with geodesic depth below 2 mm are removed (Figure 6B-6) to exclude the gyral regions from the process. Then, except for the endpoint detection step (Figure 7B), the remaining processing stages (surface filtering, surface collapse, and vertices deletion) are performed following the same approach as the one used for the gyral crowns lines estimation (Figure 6B-7–B-8). However, with respect to the endpoint detection step, there are two main differences compared to the previously described approach. First, a distance threshold of 5 mm is used to define each vertex neighborhood^27^. This condition prunes out the small branches related to possible anastomotic sulci. Second, the endpoint detection is not restricted only to vertices that form an angle lower than 30° with their corresponding neighbors. Differently from the gyral lines extraction, in this case no substantial acceleration in computation time was found when applying this restriction.

Similarly to the previous section, the result of the erosion process is a surface containing the remaining vertices and their shared faces. This mesh is then represented as a non-directed weighted graph *G*_sulc_ = (*N, E, W*), where *N* is the set of nodes (surface vertices), *E* is the set of edges connecting pairs of contiguous points, and *W* represents the edges weights (connection strength between vertices). The weight of an edge connecting vertices *i* and *j* is defined by the following mathematical expression:

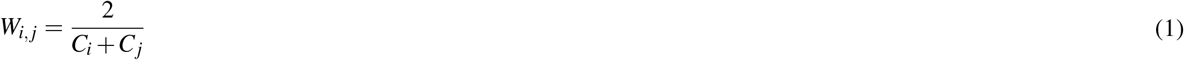

where *C*_i_ and *C*_j_ are the curvatures values for vertices *i* and *j* respectively.

A minimal spanning tree algorithm^35^ is applied on the sulcal graph with the curvature weights to find the minimal set of edges connecting all the vertices while removing possible graph cycles. Finally, the Dijkstra algorithm^36^ is applied to this graph to find the edge paths that accumulate the lowest weight value along the whole trajectory between every pair of endpoints. These paths are defined as the sulcal line corresponding to this specific basin. The output of applying this process to each sulcal basin is the set of sulcal fundi lines for a given cortical surface (see Figure 1).

### Validation

#### Sample and MRI acquisition

T1-weighted images (T1w) from the Human Connectome Project (HCP) Test-Retest dataset https://db.humanconnectome.org/^37^) were selected to evaluate the developed method. This dataset includes 45 healthy subjects (13 male, age range of 22-35 years old) scanned twice (4.7 ± 2 months interval, minimum = 1 month, maximum =11 months) on a Siemens 3T Skyra scanner in Washington University or University of Minnesota. T1w sagittal images were acquired using a Magnetization-Prepared Rapid Acquisition Gradient Echo (MPRAGE) sequence with 3D inversion recovery, echo time (TE) = 2.14 ms, repetition time (TR) = 2400 ms, inversion time (IT) = 1000 ms, flip angle (FA) = 8°, Bandwidth (BW) = 210 Hz per pixel, echo spacing (ES) = 7.6 ms, gradient strength = 42 mT/m, field of view (FOV) = 180 × 224 × 224mm^3^, voxel size = 0.7 × 0.7 × 0.7mm^3^ and an acquisition time of 7 minutes and 40 seconds. Alongside the native T1w, the HCP consortium also provides the cortical surfaces (pial and white) and their curvature maps, obtained by a project-specific FreeSurfer pipeline^38^.

#### Evaluation metrics

There are no gold standard metrics for assessing and evaluating the results obtained by the developed methodology. Previous works have used manually delineated datasets such as the MRIs Surfaces Curves (MSC) dataset^39,40^. We discarded the use of the MSC dataset because the sulcal fundi lines defined in this dataset jump, sometimes, over gyral crowns and take paths outside sulcal regions^28^. Instead, the strategy proposed for the evaluation of our method consists of comparing its results with the ones obtained by three well-known methods of sulcal features extraction: TRACE^28^, GPDM^22^, and Mindboggle^25^. All analyses are performed in ten primary sulci (five for each hemisphere) that are consistently present across individuals and show less inter-individual variability compared to other sulci: central sulcus, calcarine, superior temporal sulcus, parieto-occipital fissure, and cingulate sulcus on both hemispheres^41^. The sulcal basins for these sulci were extracted from BrainVISA’s Morphologist^21^ by projecting the sulcal median mesh over the gyral banks. These median meshes were previously manually corrected to minimize the individual differences between test and retest cortical surfaces.

##### Depth and curvature comparison

Sulcal lines should traverse the regions with higher curvature and depth. Hence, we compare ABLE against the other methods in terms of mean curvature and depth along the sulcal lines produced by any of them to see which one yields higher values of these two metrics along the lines. For this, we use the pial curvature and depth maps produced by FreeSurfer and Mindboggle, respectively. The statistical comparison of mean depth and curvature values for each sulcus is performed using paired sample t-test analyses. For each of the ten sulci, we compare ABLE with GPDM, TRACE, and Mindboggle. The resulting p-values are corrected for multiple comparisons using False Discovery Rate (FDR)^42^, considering *q* ≤ 0.05 as significant.

##### Sulcal length assessment

BrainVISA’s Morphologist pipeline^21^ has been widely used in many previous studies for the in-vivo assessment of the sulcal morphometry^5,8^. Thus, in this work, we take its length values as reference values for our comparisons. For each sulcus of interest, a linear regression analysis is performed where the length of the lines obtained by any of the methods and the reference length values are the dependent and independent variables, respectively. The p-values resulting from these regressions are corrected using FDR (*q* ≤ 0.05). Besides, sulcus-specific and global Bland-Altman plots are created to study the agreement between the sulcal lines length and the BrainVISA length values. Limits of Agreement (LoA) for Bland-Altman are estimated as the mean difference ±1.96 standard deviation.

##### Test-retest comparison

To assess the reproducibility and evaluate the spatial extraction error for each subject, we measure the mean and maximum Hausdorff distances between the lines extracted from the test and retest acquisitions using the four methods of interest. For this, we first align the retest (R) cortical surface to its correspondent test (T) cortical surface by using the Iterative Closest Point (ICP) algorithm^43^. Then, Hausdorff distances between the sulcal lines over both aligned surfaces are estimated as proposed in GPDM^22^. Concretely, we compute the Hausdorff distances between the lines obtained for the test and the aligned retest surfaces for each sulcus (see expression 2).

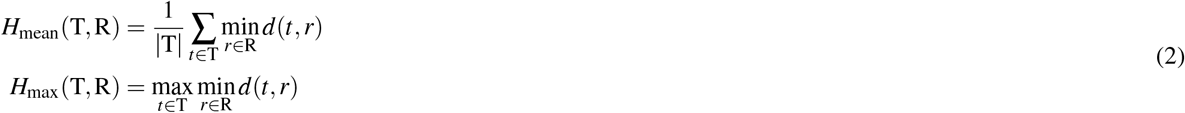

In these expressions, *d*(*t, r*) is the Euclidean distance between the vertices of the sulcal lines *t* and *r* extracted from both the test and the aligned retest cortical surfaces, respectively. *H*_mean_ reports the mean distances while *H*_max_ measures the maximum distance between the vertices on T and their closest points in R. These measures are symmetrized by averaging *H*(T, R) and *H*(R, T) for each of the metrics.

The selection of the Euclidean distance instead of geodesic distance, as proposed in GPDM^22^, responds to the spatial differences between test and retest cortical surfaces. The registration between both meshes is rigid, and therefore non-linear local deformations are not allowed. This causes that the correspondence between the vertices is not perfect and by using the Euclidean distance, we avoid projecting the vertices from one surface to the other.

The performance and reproducibility in terms of mean and maximum Hausdorff distances is evaluated using t-test analyses. As the Hausdorff distance values are not normally distributed because of the presence of outliers in this metric, the base 10 logarithm was applied to rescale the values before the statistical analysis. The p-values resulting from these tests are corrected for multiple comparisons using FDR, considering *q* ≤ 0.05 as significant.

All statistical analyses were conducted using Python’s package Pingouin^44^.

## Data availability

The datasets generated during the current study are available from the corresponding author on reasonable request.

## Acknowledgments

Data were provided by the Human Connectome Project, WU-Minn Consortium (Principal Investigators: David Van Essen and Kamil Ugurbil; 1U54MH091657) funded by the 16 NIH Institutes and Centers that support the NIH Blueprint for Neuroscience Research; and by the McDonnell Center for Systems Neuroscience at Washington University.

## Author contributions statement

Y.A.G., A.F.P., D.M.B., and J.S. conceived the method, Y.A.G., A.F.P., D.M.B., L.M.V., P.M.G, and F.J.N.S conceived the evaluation experiments and analyzed the results. A.F.P., D.M.B., S.C., J.J, Y.A.G., and M.D. worked on the redaction of the manuscript. All authors reviewed the manuscript.

