## Supplementary Material for "ABLE: Automated Brain Lines Extraction Based on Laplacian Surface Collapse"

#### Curvature and depth statistics

**Mean Curvature ($mm^{-1}$)**

| | **ABLE** | **GPDM** | **TRACE** | **Mindboggle** | **ABLE - GPDM** | | **ABLE - TRACE** | | **ABLE - Mindboggle** | |
|---|---|---|---|---|---|---|---|---|---|---|
| ***Sulcus Name*** | *Mean(sd)* | *Mean(sd)* | *Mean(sd)* | *Mean(sd)* | *$t^a$(p-FDR$^b$)* | *Cohen's D(CI$^c$)* | *t(p-FDR)* | *Cohen's D(CI)* | *t(p-FDR)* | *Cohen's D(CI)* |
| *Left Hemisphere* | | | | | | | | | | |
| Calcarine | 0.348 (0.058) | 0.364 (0.101) | 0.259 (0.042) | 0.168 (0.101) | -2.19 (0.255) | -0.30 (-0.60, -0.01) | 18.88 (**<0.001**) | 1.93 (1.44, 2.43) | 13.60 (**<0.001**) | 2.44 (1.86, 3.03) |
| Central Sulcus | 0.436 (0.048) | 0.429 (0.059) | 0.309 (0.032) | 0.326 (0.046) | 1.38 (1.000) | 0.15 (-0.14, 0.45) | 24.40 (**<0.001**) | 3.63 (2.82, 4.43) | 21.79 (**<0.001**) | 2.73 (2.10, 3.37) |
| Cingulate | 0.327 (0.047) | 0.294 (0.090) | 0.256 (0.034) | 0.293 (0.040) | 2.92 (0.051) | 0.58 (0.26, 0.90) | 17.15 (**<0.001**) | 2.21 (1.67, 2.75) | 6.85 (**<0.001**) | 0.97 (0.62, 1.33) |
| Parieto-Occipital Fissure | 0.303 (0.075) | 0.346 (0.130) | 0.261 (0.031) | 0.263 (0.059) | -2.58 (0.110) | -0.51 (-0.82, -0.20) | 4.84 (**<0.001**) | 0.94 (0.59, 1.29) | 4.43 (**0.001**) | 0.76 (0.43, 1.09) |
| Superior Temporal | 0.307 (0.056) | 0.298 (0.077) | 0.221 (0.035) | 0.237 (0.051) | 1.53 (0.912) | 0.16 (-0.14, 0.45) | 19.18 (**<0.001**) | 2.13 (1.60, 2.66) | 11.38 (**<0.001**) | 1.49 (1.06, 1.91) |
| *Right Hemisphere* | | | | | | | | | | |
| Calcarine | 0.354 (0.065) | 0.395 (0.096) | 0.272 (0.031) | 0.192 (0.067) | -4.06 (**0.002**) | -0.61 (-0.92, -0.29) | 11.91 (**<0.001**) | 1.98 (1.48, 2.49) | 14.22 (**<0.001**) | 2.91 (2.24, 3.58) |
| Central Sulcus | 0.401 (0.045) | 0.400 (0.050) | 0.278 (0.023) | 0.304 (0.037) | 0.26 (1.000) | 0.03 (-0.26, 0.32) | 25.63 (**<0.001**) | 4.25 (3.33, 5.18) | 18.10 (**<0.001**) | 2.73 (2.09, 3.36) |
| Cingulate | 0.336 (0.044) | 0.272 (0.079) | 0.258 (0.028) | 0.290 (0.037) | 6.57 (**<0.001**) | 1.08 (0.71, 1.45) | 18.93 (**<0.001**) | 2.40 (1.83, 2.98) | 12.42 (**<0.001**) | 1.22 (0.83, 1.60) |
| Parieto-Occipital Fissure | 0.295 (0.055) | 0.341 (0.125) | 0.259 (0.036) | 0.259 (0.052) | -3.12 (**0.031**) | -0.66 (-0.98, -0.34) | 5.67 (**<0.001**) | 0.92 (0.57, 1.27) | 5.32 (**<0.001**) | 0.80 (0.47, 1.14) |
| Superior Temporal | 0.305 (0.048) | 0.293 (0.078) | 0.226 (0.108) | 0.238 (0.047) | 1.59 (0.829) | 0.23 (-0.06, 0.53) | 6.83 (**<0.001**) | 1.33 (0.93, 1.73) | 10.14 (**<0.001**) | 1.83 (1.36, 2.31) |

**Mean Depth (mm)**

| | **ABLE** | **GPDM** | **TRACE** | **Mindboggle** | **ABLE - GPDM** | | **ABLE - TRACE** | | **ABLE - Mindboggle** | |
|---|---|---|---|---|---|---|---|---|---|---|
| ***Sulcus Name*** | *Mean(sd)* | *Mean(sd)* | *Mean(sd)* | *Mean(sd)* | *$t^a$(p-FDR$^b$)* | *Cohen's D(CI$^c$)* | *t(p-FDR)* | *Cohen's D(CI)* | *t(p-FDR)* | *Cohen's D(CI)* |
| *Left Hemisphere* | | | | | | | | | | |
| Calcarine | 12.48 (1.55) | 15.19 (3.06) | 11.80 (1.49) | 11.73 (2.77) | -7.77 (**<0.001**) | -1.28 (-1.67, -0.88) | 5.50 (**<0.001**) | 0.45 (0.14, 0.75) | 1.85 (0.513) | 0.36 (0.06, 0.66) |
| Central Sulcus | 14.27 (1.18) | 17.27 (1.48) | 12.27 (0.90) | 13.05 (0.95) | -26.19 (**<0.001**) | -2.31 (-2.87, -1.75) | 21.34 (**<0.001**) | 1.98 (1.48, 2.48) | 12.34 (**<0.001**) | 1.20 (0.82, 1.59) |
| Cingulate | 9.16 (1.13) | 8.27 (1.45) | 8.99 (0.81) | 9.73 (0.84) | 4.25 (**0.001**) | 0.73 (0.40, 1.06) | 1.86 (0.512) | 0.19 (-0.11, 0.48) | -4.70 (**<0.001**) | -0.62 (-0.94, -0.30) |
| Parieto-Occipital Fissure | 13.59 (1.78) | 16.69 (3.12) | 13.46 (1.54) | 14.25 (1.95) | -8.33 (**<0.001**) | -1.40 (-1.81, -0.99) | 0.59 (1.000) | 0.07 (-0.22, 0.36) | -2.78 (0.068) | -0.40 (-0.71, -0.10) |
| Superior Temporal | 14.02 (1.80) | 15.81 (2.43) | 12.66 (1.51) | 12.58 (1.49) | -10.46 (**<0.001**) | -0.86 (-1.20, -0.52) | 12.61 (**<0.001**) | 0.83 (0.49, 1.17) | 8.54 (**<0.001**) | 0.90 (0.55, 1.25) |
| *Right Hemisphere* | | | | | | | | | | |
| Calcarine | 13.03 (1.62) | 16.24 (2.71) | 12.37 (1.74) | 12.72 (2.56) | -11.50 (**<0.001**) | -1.64 (-2.08, -1.19) | 3.96 (**0.003**) | 0.41 (0.10, 0.71) | 1.24 (1.000) | 0.20 (-0.10, 0.49) |
| Central Sulcus | 14.47 (1.07) | 17.69 (1.24) | 12.75 (0.93) | 13.32 (0.85) | -27.55 (**<0.001**) | -2.86 (-3.52, -2.20) | 14.36 (**<0.001**) | 1.77 (1.30, 2.24) | 9.46 (**<0.001**) | 1.26 (0.87, 1.65) |
| Cingulate | 9.58 (1.18) | 9.00 (1.82) | 9.21 (0.98) | 9.60 (0.84) | 3.24 (**0.023**) | 0.43 (0.12, 0.73) | 3.45 (**0.013**) | 0.30 (0.00, 0.60) | -0.34 (1.000) | -0.03 (-0.32, 0.26) |
| Parieto-Occipital Fissure | 14.59 (1.53) | 18.43 (3.19) | 14.59 (1.54) | 15.50 (2.03) | -9.45 (**<0.001**) | -1.68 (-2.13, -1.23) | -0.06 (1.000) | -0.01 (-0.30, 0.28) | -4.42 (**<0.001**) | -0.54 (-0.86, -0.23) |
| Superior Temporal | 15.85 (1.79) | 19.22 (2.63) | 14.10 (1.23) | 14.20 (1.63) | -14.34 (**<0.001**) | -1.52 (-1.95, -1.09) | 14.26 (**<0.001**) | 1.19 (0.81, 1.57) | 9.58 (**<0.001**) | 1.00 (0.64, 1.36) |

[a] *Comparisons between groups were conducted using the pairwise t-test.*
[b] *P-values in bold remain significant after entering all uncorrected p-values (N = 120) into a False Discovery Rate (FDR) correction for multiple comparisons with q = 0.05.*
[c] *CI≡ 95% confidence interval.*

**Table S1.** Comparison of the values of mean sulcal fundi curvatures and depths. p-values FDR corrected.

### Hausdorff distances statistics

##### *log* (Mean Hausdorff Distances)

| | ABLE | GPDM | TRACE | Mindboggle | ABLE - GPDM | | ABLE - TRACE | | ABLE - Mindboggle | |
|---|---|---|---|---|---|---|---|---|---|---|
| ***Sulcus Name*** | *Mean(sd)* | *Mean(sd)* | *Mean(sd)* | *Mean(sd)* | *t[a](p-FDR[b])* | *Cohen's D(CI[c])* | *t(p-FDR)* | *Cohen's D(CI)* | *t(p-FDR)* | *Cohen's D(CI)* |
| *Left Hemisphere* | | | | | | | | | | |
| Calcarine | 0.08 (0.18) | 0.19 (0.33) | -0.07 (0.08) | 0.27 (0.20) | -1.73 (0.627) | -0.34 (-0.79, 0.11) | 5.55 (**<0.001**) | 1.0703 (0.52, 1.62) | -4.81 (**<0.001**) | -0.96 (-1.49, -0.43) |
| Central Sulcus | -0.18 (0.19) | -0.23 (0.21) | -0.16 (0.09) | 0.12 (0.20) | 1.25 (1.000) | 0.26 (-0.18, 0.71) | -0.62 (1.000) | -0.1338 (-0.57, 0.31) | -7.40 (**<0.001**) | -1.52 (-2.16, -0.88) |
| Cingulate | 0.01 (0.15) | 0.10 (0.41) | -0.08 (0.08) | 0.02 (0.15) | -1.29 (1.000) | -0.26 (-0.71, 0.18) | 4.38 (**0.001**) | 0.7928 (0.29, 1.29) | 0.58 (1.000) | 0.13 (-0.31, 0.57) |
| Parieto-Occipital Fissure | 0.10 (0.26) | 0.39 (0.40) | -0.04 (0.15) | 0.16 (0.20) | -4.04 (**0.003**) | -0.86 (-1.37, -0.34) | 3.50 (**0.012**) | 0.7010 (0.21, 1.19) | -0.97 (1.000) | -0.23 (-0.67, 0.22) |
| Superior Temporal | -0.07 (0.16) | -0.05 (0.27) | -0.12 (0.09) | 0.15 (0.18) | -0.29 (1.000) | -0.07 (-0.50, 0.37) | 2.74 (0.074) | 0.4610 (0.00, 0.92) | -5.66 (**<0.001**) | -1.28 (-1.87, -0.69) |
| *Right Hemisphere* | | | | | | | | | | |
| Calcarine | 0.07 (0.18) | 0.21 (0.36) | -0.06 (0.10) | 0.26 (0.15) | -2.10 (0.314) | -0.41 (-0.87, 0.04) | 5.11 (**<0.001**) | 0.9917 (0.46, 1.53) | -4.63 (**<0.001**) | -1.07 (-1.62, -0.52) |
| Central Sulcus | -0.17 (0.19) | -0.25 (0.18) | -0.16 (0.07) | 0.07 (0.17) | 1.86 (0.490) | 0.41 (-0.05, 0.86) | -0.48 (1.000) | -0.0951 (-0.53, 0.34) | -6.37 (**<0.001**) | -1.33 (-1.94, -0.73) |
| Cingulate | -0.02 (0.18) | 0.08 (0.34) | -0.08 (0.12) | 0.00 (0.19) | -1.65 (0.732) | -0.39 (-0.84, 0.07) | 2.01 (0.380) | 0.3050 (-0.14, 0.75) | -0.82 (1.000) | -0.16 (-0.60, 0.28) |
| Parieto-Occipital Fissure | 0.09 (0.24) | 0.10 (0.36) | -0.07 (0.13) | 0.17 (0.22) | -0.13 (1.000) | -0.03 (-0.47, 0.41) | 4.61 (**0.001**) | 0.8690 (0.36, 1.38) | -1.92 (0.450) | -0.37 (-0.82, 0.08) |
| Superior Temporal | 0.08 (0.19) | -0.03 (0.27) | -0.12 (0.06) | 0.19 (0.17) | 2.37 (0.177) | 0.52 (0.05, 0.99) | 7.30 (**<0.001**) | 1.4781 (0.84, 2.11) | -2.61 (0.102) | -0.56 (-1.03, -0.09) |

##### *log* (Maximum Hausdorff Distances)

| | ABLE | GPDM | TRACE | Mindboggle | ABLE - GPDM | | ABLE - TRACE | | ABLE - Mindboggle | |
|---|---|---|---|---|---|---|---|---|---|---|
| ***Sulcus Name*** | *Mean(sd)* | *Mean(sd)* | *Mean(sd)* | *Mean(sd)* | *t[a](p-FDR[b])* | *Cohen's D(CI[c])* | *t(p-FDR)* | *Cohen's D(CI)* | *t(p-FDR)* | *Cohen's D(CI)* |
| *Left Hemisphere* | | | | | | | | | | |
| Calcarine | 0.87 (0.15) | 0.78 (0.27) | 0.81 (0.10) | 0.98 (0.25) | 2.20 (0.257) | 0.44 (-0.02, 0.90) | 3.42 (**0.014**) | 0.55 (0.08, 1.02) | -2.50 (0.131) | -0.51 (-0.9700, -0.04) |
| Central Sulcus | 0.71 (0.29) | 0.49 (0.28) | 0.85 (0.14) | 1.00 (0.16) | 3.81 (**0.004**) | 0.80 (0.30, 1.30) | -2.70 (0.081) | -0.59 (-1.07, -0.12) | -6.12 (**<0.001**) | -1.21 (-1.7800, -0.63) |
| Cingulate | 0.90 (0.18) | 0.70 (0.44) | 0.93 (0.09) | 0.95 (0.15) | 2.86 (0.057) | 0.61 (0.14, 1.09) | -0.62 (1.000) | -0.12 (-0.56, 0.32) | -0.93 (1.000) | -0.20 (-0.6400, 0.24) |
| Parieto-Occipital Fissure | 0.80 (0.25) | 0.82 (0.33) | 0.83 (0.17) | 0.93 (0.15) | -0.35 (1.000) | -0.07 (-0.50, 0.37) | -0.67 (1.000) | -0.12 (-0.56, 0.32) | -3.05 (**0.036**) | -0.63 (-1.1100, -0.15) |
| Superior Temporal | 0.84 (0.22) | 0.67 (0.28) | 0.87 (0.13) | 1.01 (0.15) | 3.14 (**0.028**) | 0.69 (0.20, 1.18) | -0.75 (1.000) | -0.13 (-0.57, 0.31) | -4.26 (**0.001**) | -0.90 (-1.4200, -0.38) |
| *Right Hemisphere* | | | | | | | | | | |
| Calcarine | 0.88 (0.18) | 0.78 (0.28) | 0.84 (0.11) | 1.02 (0.15) | 2.61 (0.103) | 0.53 (0.07, 1.00) | 2.29 (0.211) | 0.46 (-0.00, 0.92) | -4.18 (**0.002**) | -0.79 (-1.2900, -0.29) |
| Central Sulcus | 0.64 (0.31) | 0.44 (0.25) | 0.83 (0.11) | 0.97 (0.14) | 3.36 (**0.016**) | 0.70 (0.21, 1.19) | -4.33 (**0.001**) | -0.82 (-1.32, -0.31) | -7.18 (**<0.001**) | -1.37 (-1.9800, -0.76) |
| Cingulate | 0.93 (0.20) | 0.68 (0.40) | 0.92 (0.15) | 0.96 (0.17) | 3.04 (**0.038**) | 0.72 (0.23, 1.21) | -0.37 (1.000) | -0.07 (-0.50, 0.37) | -1.16 (1.000) | -0.25 (-0.7000, 0.19) |
| Parieto-Occipital Fissure | 0.82 (0.21) | 0.59 (0.33) | 0.77 (0.16) | 0.92 (0.18) | 3.80 (**0.005**) | 0.85 (0.34, 1.36) | 1.57 (0.827) | 0.32 (-0.13, 0.76) | -2.09 (0.320) | -0.46 (-0.9200, 0.00) |
| Superior Temporal | 0.92 (0.17) | 0.65 (0.27) | 0.86 (0.09) | 1.04 (0.16) | 5.22 (**<0.001**) | 1.19 (0.61, 1.76) | 1.87 (0.490) | 0.39 (-0.06, 0.85) | -3.86 (**0.004**) | -0.75 (-1.2400, -0.25) |

[a] *Comparisons between groups were conducted using the pairwise t-test.*
[b] *P-values in bold remain significant after entering all uncorrected p-values (N = 120) into a False Discovery Rate (FDR) correction for multiple comparisons with q = 0.05.*
[c] *CI≡95% confidence interval.*

**Table S2.** Comparison of the values of the logarithms of mean Hausdorff and maximum Hausdorff distances yielded by the different methods. Symbols "$*$", "$**$", and "$***$", stand for significant differences from a t-test with FDR-corrected p-values lower than 0.05, 0.01, and 0.001, respectively.

### Bland-Altman plots per sulcus

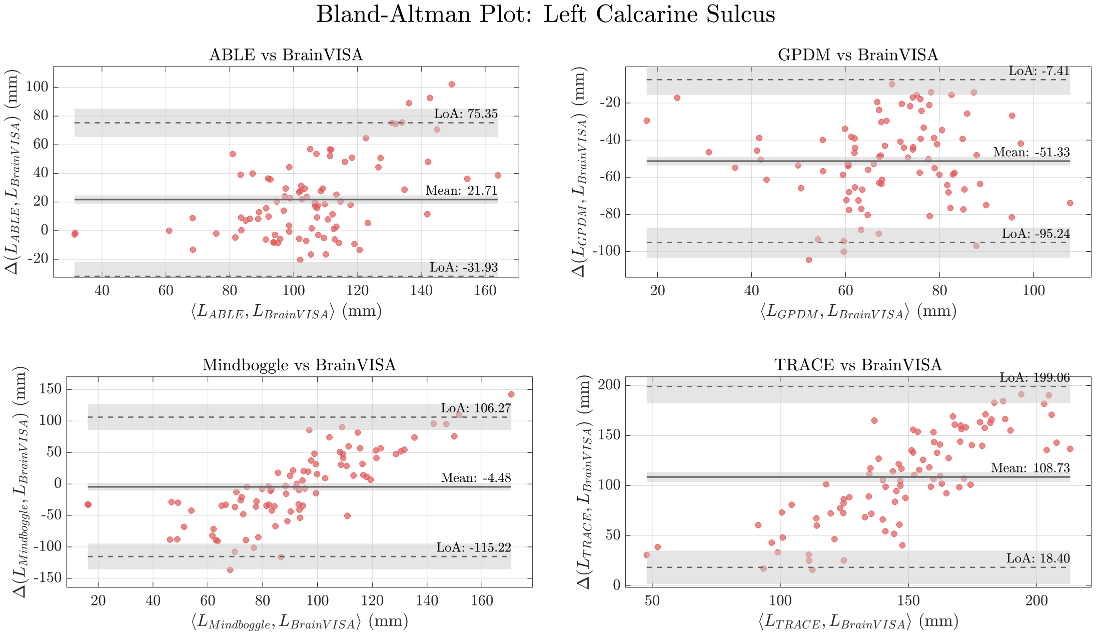


**Figure S1.** Bland-Altman plot comparing the four methods (ABLE, GPDM, Mindboggle, and TRACE) against BrainVISA's gold standard for the left calcarine sulcus.

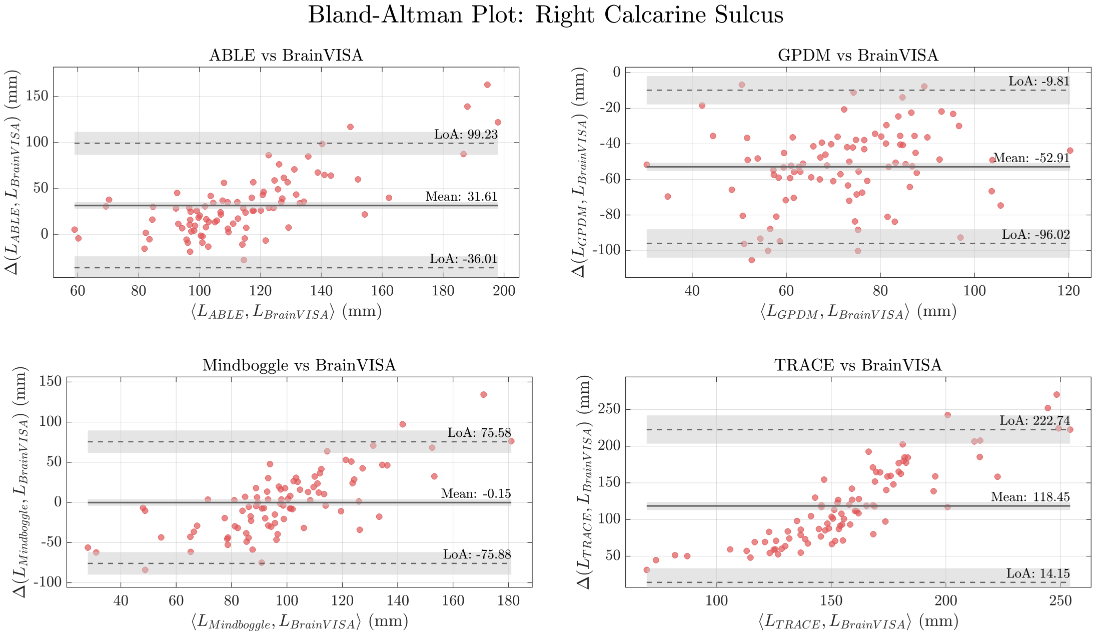


**Figure S2.** Bland-Altman plot comparing the four methods (ABLE, GPDM, Mindboggle, and TRACE) against BrainVISA's gold standard for the right calcarine sulcus.

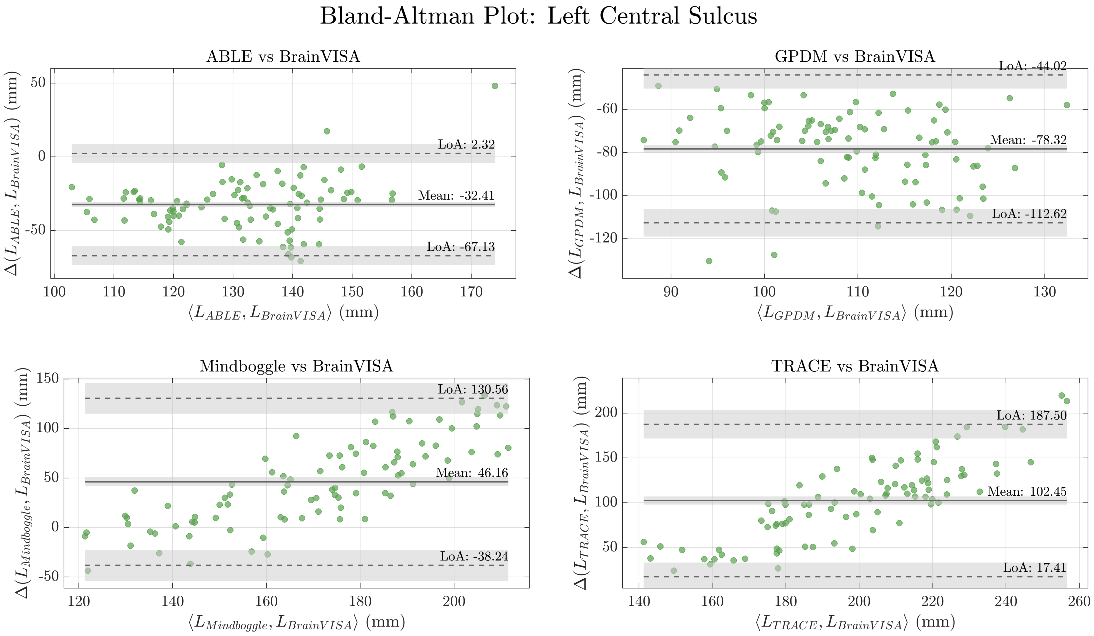


**Figure S3.** Bland-Altman plot comparing the four methods (ABLE, GPDM, Mindboggle, and TRACE) against BrainVISA's gold standard for the left central sulcus.

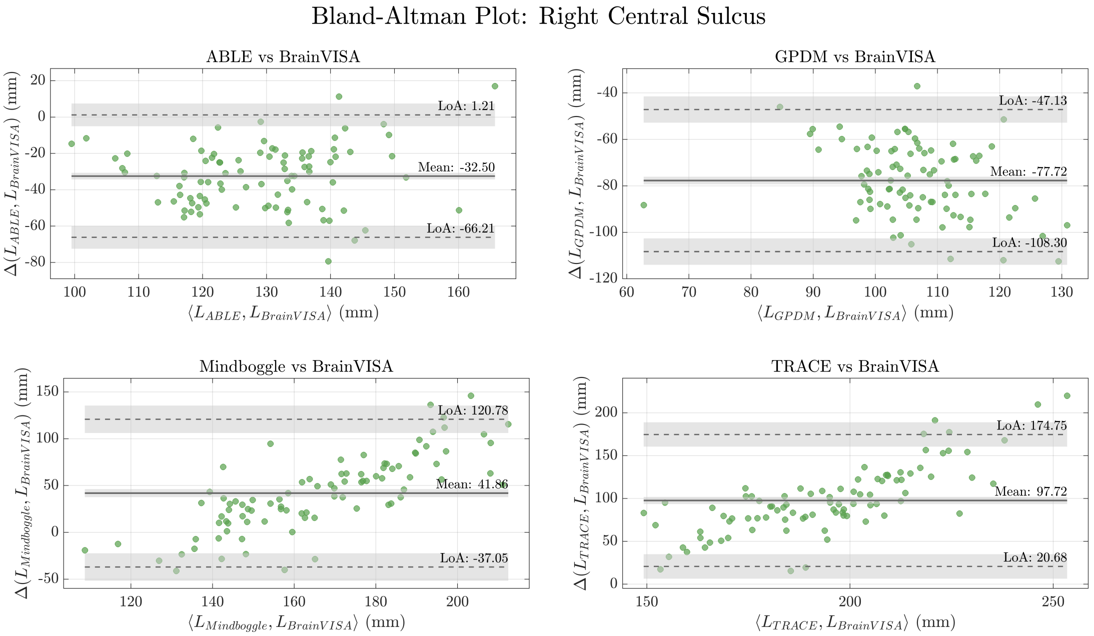


**Figure S4.** Bland-Altman plot comparing the four methods (ABLE, GPDM, Mindboggle, and TRACE) against BrainVISA's gold standard for the right central sulcus.

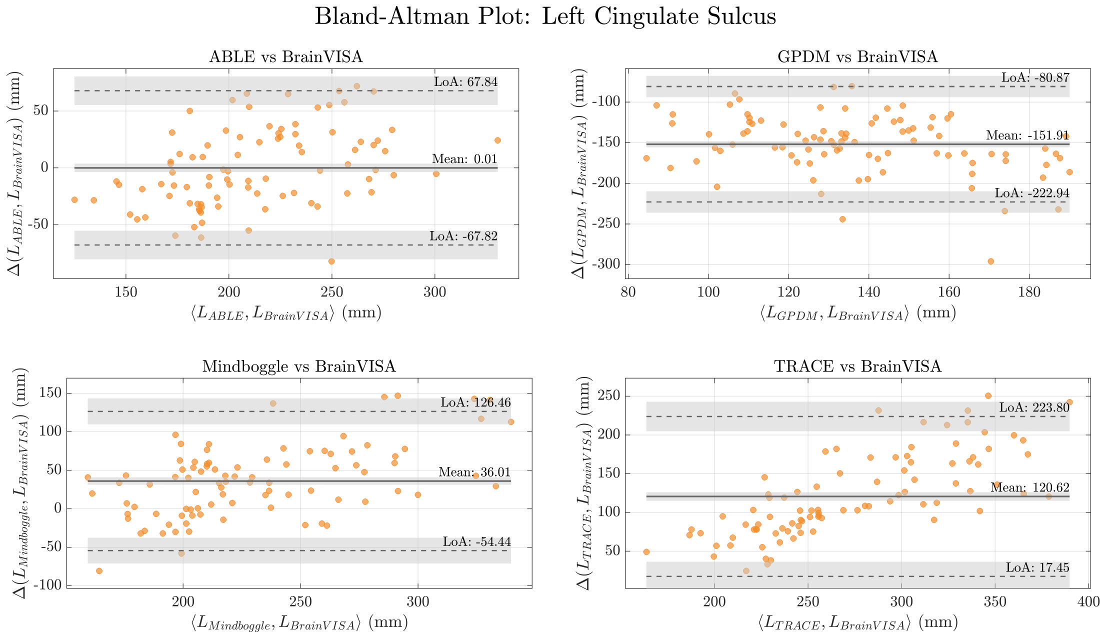


**Figure S5.** Bland-Altman plot comparing the four methods (ABLE, GPDM, Mindboggle, and TRACE) against BrainVISA's gold standard for the left cingulate sulcus.

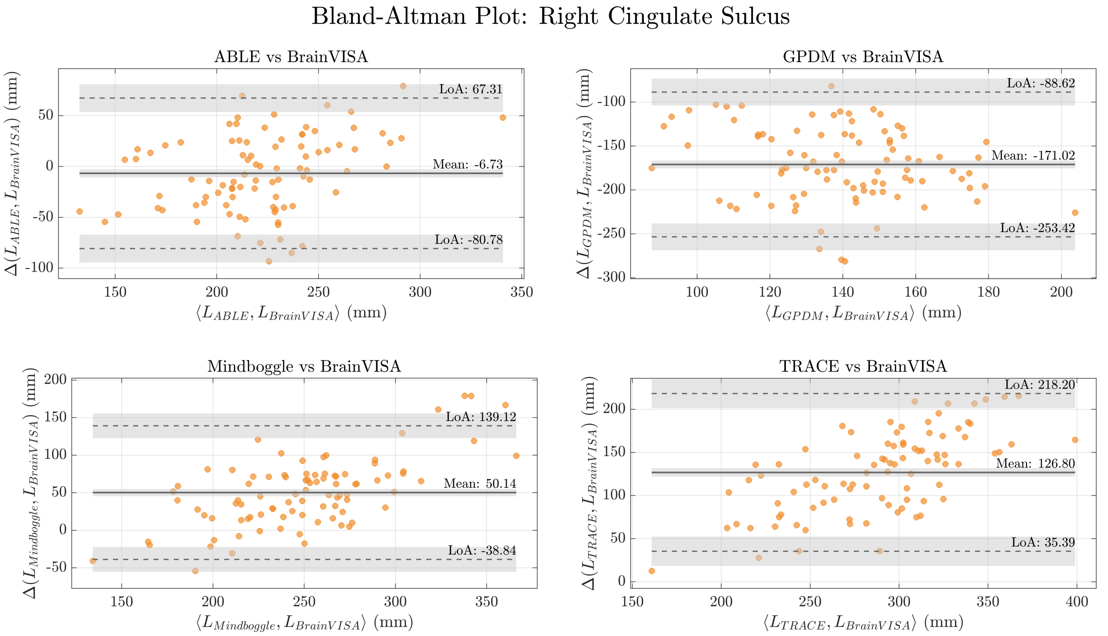


**Figure S6.** Bland-Altman plot comparing the four methods (ABLE, GPDM, Mindboggle, and TRACE) against BrainVISA's gold standard for the right cingulate sulcus.

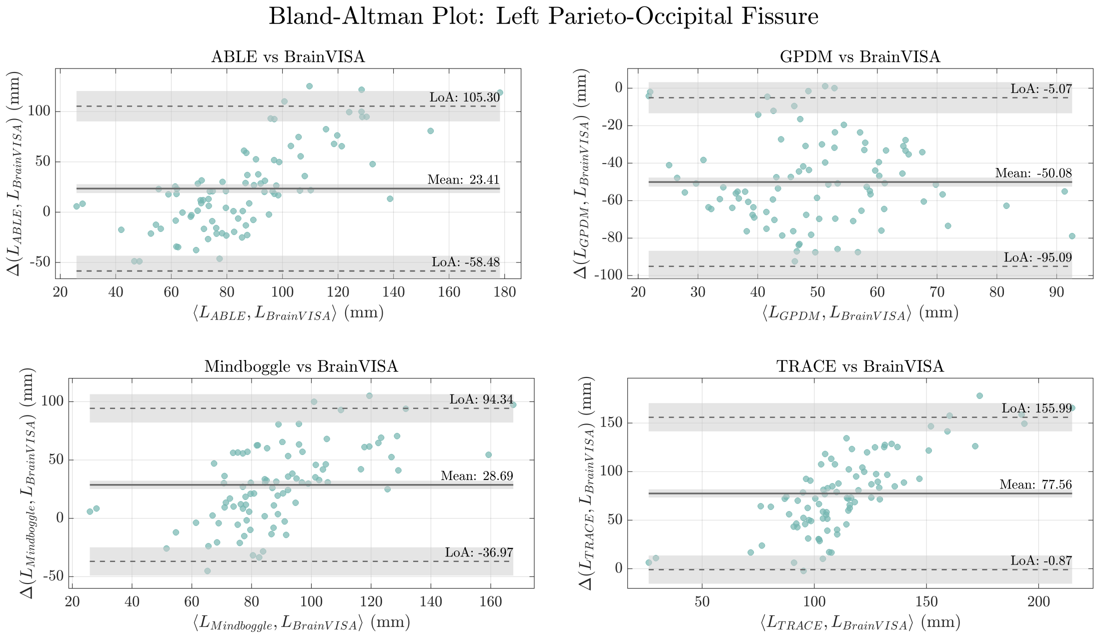


**Figure S7.** Bland-Altman plot comparing the four methods (ABLE, GPDM, Mindboggle, and TRACE) against BrainVISA's gold standard for the left parieto-occipital fissure.

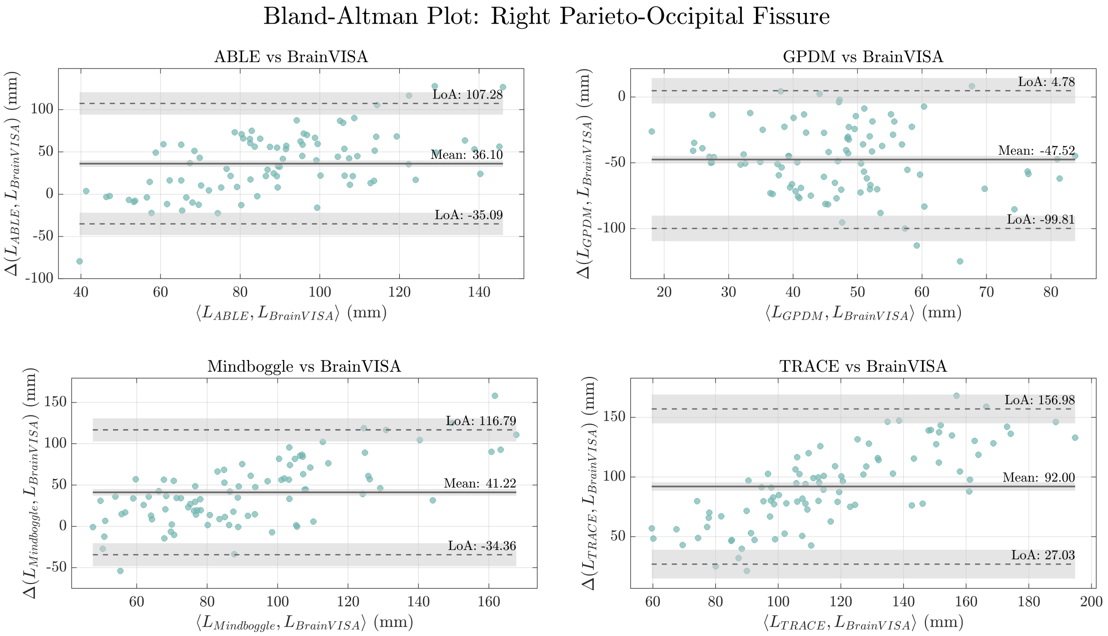


**Figure S8.** Bland-Altman plot comparing the four methods (ABLE, GPDM, Mindboggle, and TRACE) against BrainVISA's gold standard for the right parieto-occipital fissure.

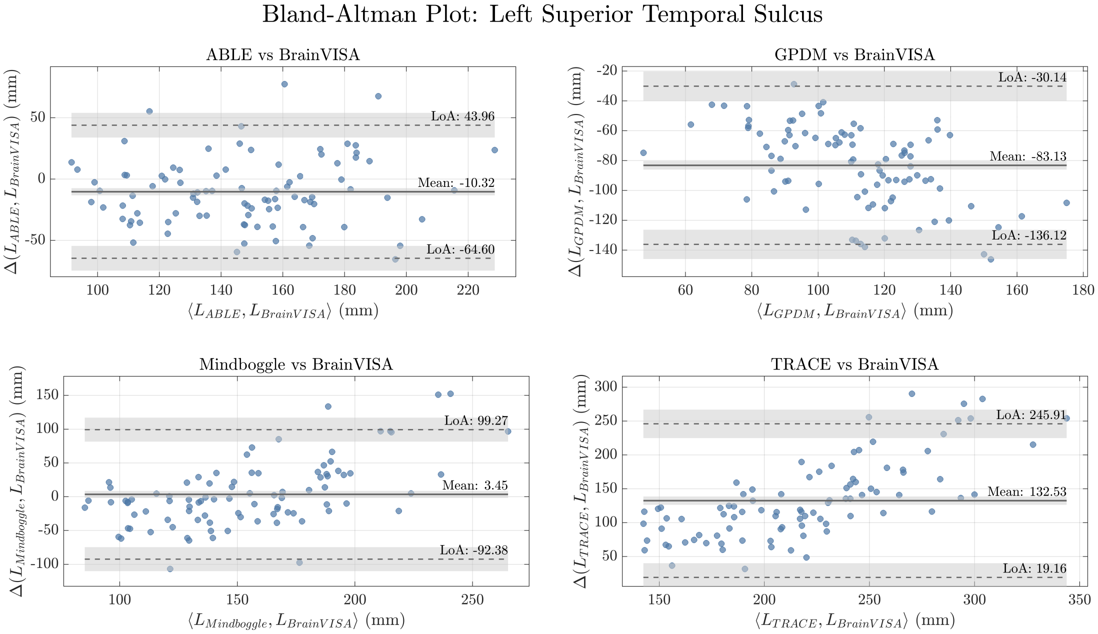


**Figure S9.** Bland-Altman plot comparing the four methods (ABLE, GPDM, Mindboggle, and TRACE) against BrainVISA's gold standard for the left superior temporal sulcus.

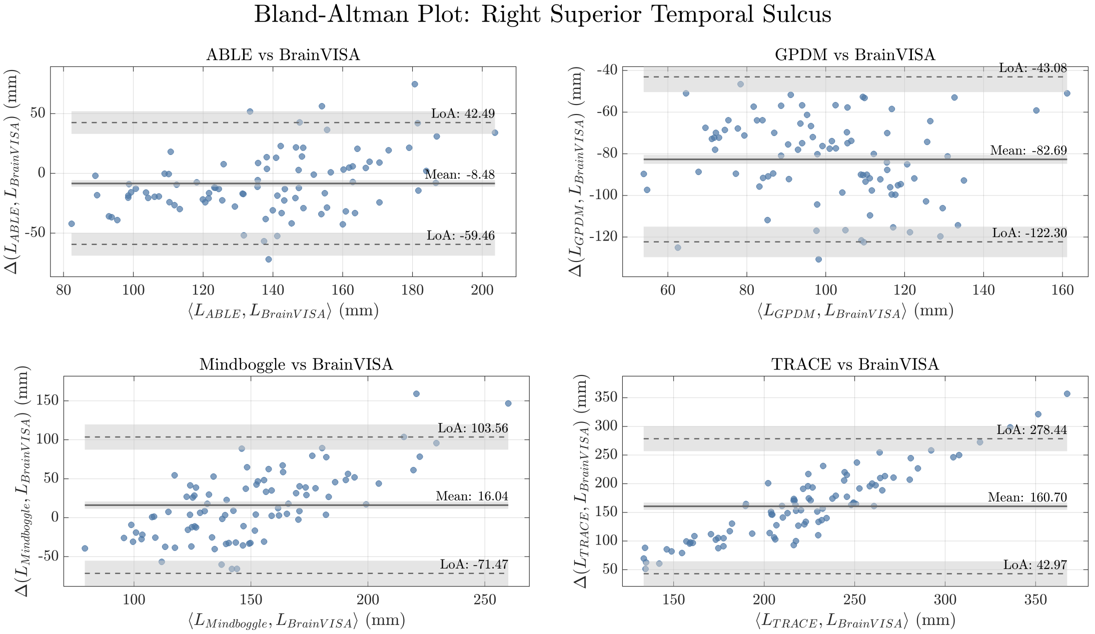


**Figure S10.** Bland-Altman plot comparing the four methods (ABLE, GPDM, Mindboggle, and TRACE) against BrainVISA's gold standard for the right superior temporal sulcus.

#### Summary of longitude differences between methods and reference values

**Differences between methods and BrainVISA (mm)**

| ***Sulcus Name*** | **ABLE**<br>*Mean (CI[2.5%], CI[97.5%])* | **GPDM**<br>*Mean (CI[2.5%], CI[97.5%])* | **TRACE**<br>*Mean (CI[2.5%], CI[97.5%])* | **Mindboggle**<br>*Mean (CI[2.5%], CI[97.5%])* |
|---|---|---|---|---|
| *Left Hemisphere* | | | | |
| *Calcarine* | 21.71 (-31.94, 75.35) | -51.33 (-95.24, -7.41) | 108.73 (18.40, 199.06) | -4.48 (-115.22, 106.27) |
| *Central Sulcus* | -32.41 (-67.13, 2.32) | -78.32 (-112.62, -44.02) | 102.45 (17.41, 187.50) | 46.16 (-38.24, 130.56) |
| *Cingulate* | 0.01 (-67.82, 67.84) | -151.91 (-222.94, -80.87) | 120.62 (17.45, 223.80) | 36.01 (-54.44, 126.46) |
| *Parieto-Occipital Fissure* | 23.41 (-58.48, 105.31) | -50.08 (-95.09, -5.07) | 77.56 (-0.87, 155.99) | 28.69 (-36.97, 94.34) |
| *Superior Temporal* | -10.32 (-64.60, 43.96) | -83.13 (-136.12, -30.14) | 132.53 (19.16, 245.91) | 3.45 (-92.38, 99.27) |
| *Right Hemisphere* | | | | |
| *Calcarine* | 31.61 (-36.01, 99.23) | -52.91 (-96.02, -9.81) | 118.45 (14.15, 222.75) | -0.15 (-75.88, 75.58) |
| *Central Sulcus* | -32.50 (-66.21, 1.21) | -77.72 (-108.30, -47.13) | 97.72 (20.68, 174.75) | 41.86 (-37.05, 120.78) |
| *Cingulate* | -6.73 (-80.78, 67.31) | -171.02 (-253.42, -88.62) | 126.80 (35.39, 218.21) | 50.14 (-38.85, 139.12) |
| *Parieto-Occipital Fissure* | 36.10 (-35.09, 107.28) | -47.52 (-99.81, 4.78) | 92.00 (27.03, 156.98) | 41.22 (-34.36, 116.79) |
| *Superior Temporal* | -8.48 (-59.46, 42.49) | -82.69 (-122.30, -43.08) | 160.70 (42.97, 278.44) | 16.04 (-71.47, 103.56) |

**Table S3.** Summary of mean differences between methods and confidence intervals extracted from the Bland-Altman plots.

#### Intercept of sulcal longitudes regressions

| Sulcus Name | ABLE<br>Intercept CI([2.5%],[97.5%]) | GPDM<br>Intercept CI([2.5%],[97.5%]) | TRACE<br>Intercept CI([2.5%],[97.5%]) | Mindboggle<br>Intercept CI([2.5%],[97.5%]) |
|---|---|---|---|---|
| *Left Hemisphere* | | | | |
| *Calcarine* | 38.68 (9.67, 67.68) | 11.87 (-7.79, 31.52) | 71.24 (22.68, 119.80) | 99.59 (43.58, 155.60) |
| *Central Sulcus* | 42.96 (7.33, 78.60) | 55.57 (29.52, 81.61) | 94.39 (-1.31, 190.09) | 105.62 (11.48, 199.76) |
| *Cingulate* | 5.34 (-37.44, 48.12) | 0.64 (-29.90, 31.18) | 12.27 (-48.51, 73.06) | 34.66 (-22.40, 91.73) |
| *Parieto-Occipital Fissure* | 44.76 (8.10, 81.43) | 13.16 (-1.75, 28.07) | 61.23 (26.01, 96.45) | 52.52 (23.35, 81.69) |
| *Superior Temporal* | 38.17 (11.64, 64.71) | 18.90 (1.91, 35.89) | 91.67 (32.73, 150.62) | 15.58 (-34.73, 65.89) |
| *Right Hemisphere* | | | | |
| *Calcarine* | 11.97 (-29.59, 53.54) | 0.03 (-24.04, 24.10) | 48.42 (-14.23, 111.07) | -0.92 (-47.71, 45.86) |
| *Central Sulcus* | 57.95 (26.47, 89.43) | 41.89 (20.15, 63.63) | 154.23 (70.76, 237.70) | 102.18 (16.75, 187.61) |
| *Cingulate* | 43.76 (-9.38, 96.91) | 44.87 (6.09, 83.66) | 75.36 (9.32, 141.39) | 0.06 (-64.22, 64.35) |
| *Parieto-Occipital Fissure* | 61.29 (33.91, 88.67) | 23.77 (10.56, 36.99) | 53.19 (29.17, 77.22) | 41.74 (12.08, 71.39) |
| *Superior Temporal* | 3.69 (-28.44, 35.82) | -20.54 (-41.73, 0.65) | -36.53 (-97.74, 24.67) | 8.66 (-46.66, 63.98) |

**Table S4.** Intercept (and its confidence interval) of the linear regressions of the sulcal longitudes obtained by each of the methods and gold standard.
